# Screening for variable drug responses using human iPSC cohorts

**DOI:** 10.1101/2023.06.16.545161

**Authors:** Melpomeni Platani, Hao Jiang, Lindsay Davidson, Santosh Hariharan, Regis Doyonnas, Angus I. Lamond, Jason R. Swedlow

## Abstract

We have developed a laboratory-based drug screening platform that uses a cohort of human induced pluripotent stem cell (hiPSC) lines, derived from different donors, to predict variable drug responses of potential clinical relevance. This builds on recent findings that pluripotent hiPSC lines express a broad repertoire of gene transcripts and proteins, whose expression levels reflect the genetic identity of the donor. We demonstrate that a cohort of hiPSC lines from different donors can be screened efficiently in their pluripotent state, using high-throughput Cell Painting assays. Variable phenotypic responses between hiPSC lines were detected with a wide range of clinically approved drugs, in use across multiple disease areas. Furthermore, information on mechanisms of drug-cell interactions underlying the observed variable responses was derived by using quantitative proteomic analysis to compare sets of hiPSC lines that had been stratified objectively, based upon variable response, Cell Painting data. We propose that information derived from comparative drug screening, using curated libraries of hiPSC lines from different donors, can help to improve the delivery of safe new drugs suitable for a broad range of genetic backgrounds and sexual diversity within human populations.

## INTRODUCTION

Most current drug development pipelines involve an initial laboratory-based research phase, in which disease-relevant molecular targets are identified (usually proteins) and then either small molecules, or biological effectors (e.g. antibodies), are characterised that specifically bind to, or otherwise modulate, the targets. Promising drug candidates are then progressed from animal models to human clinical trials, where they must be evaluated for efficacy and possible toxicity in human patients before they can be certified for use by regulatory authorities (*1–3*). Unfortunately, these pipelines often have very high failure rates, with 80% or more drug candidates in major disease areas failing to pass clinical trials (*4*). The high failure rates at late stages of drug development are a major factor in the high cost of bringing safe and effective new drugs to market.

Another significant limitation with current pre-clinical drug development pipelines is the failure to account for the suitability of drugs for use with diverse patient groups from different sexes and/or genetic backgrounds. Variable drug responses resulting from natural genetic variations in human populations is recognised to contribute to the high failure rates of drug candidates in clinical trials (*5*). Fundamentally, drug development pipelines that focus on mapping drug molecules to targets in the laboratory, do not account for normal population-level differences between individuals, which affect the outcome of clinical trials on patients.

There are now many examples of drugs, used in oncology (*6*), anti-psychotics (*7*), cholesterol-lowering medications (*8*) and others (*9*), where variable efficacy and toxicity are well-documented and included in standard dosing guidelines. In many cases, differential responses result from epistatic interactions and have been mapped to genetic variants affecting the expression levels and/or activity of cell proteins and pathways, for example, metabolic enzymes, membrane transporters and signalling molecules etc. A key gap in the drug development pipeline, therefore, is the lack of assays that report on the impact of population-level phenotypic variation during the pre-clinical phases of drug development.

Several projects have now established cohorts of human induced pluripotent cell (hiPSC) lines from multiple donors using standardised protocols (*10, 11*). In particular, the Human Induced Pluripotent Stem Cell Initiative, (HipSci, https://www.hipsci.org), has established a library of >700 cell lines, generated from over 300 individuals, including healthy male and female donors of varied ages, as well as patient cohorts with known genetic disorders (*12*). The HipSci cell lines were all reprogrammed from human skin fibroblasts, following transduction of the pluripotency transcription factors using a Sendai virus vector (*12*). Many of these cell lines have been subjected to mRNA and protein expression analysis and these data are publicly available. Interestingly, comparison of mRNA and protein expression patterns in the HipSci lines showed that mRNA and protein levels in separate hiPSC lines from the same individual are more similar to each other than to lines derived from different donors. This shows that gene expression states of the HipSci lines are strongly defined by the genetic identity of the donor (*12, 13*). The HipSci cohort therefore provides a tractable system for measuring how human genetic variation affects cellular phenotypes, potentially including variable drug responses.

The ability of hiPSCs to be re-differentiated into different cell types, using standardised differentiation protocols, is frequently used for experimental analyses of tissue-specific responses and disease mechanisms (*14*). However, using the undifferentiated, pluripotent state of hiPSCs for drug screening provides some potential advantages. For example, studies that have measured in depth gene transcript and protein expression in hiPSCs and embryonic stem cells (ESCs) (*13, 15*), find that stem cells express a broader repertoire of gene transcripts and proteins than terminally differentiated, primary cells and tissues, including most tumour-derived, transformed cell lines. For example, hiPSC lines express ∼2.5-fold more receptor tyrosine kinases than many primary cells, or tumour cell lines. For the majority of chromosomes, >50% of their known protein coding genes are expressed in hiPSCs ((*13*) and references therein). While no single cell type expresses all potential drug targets, this broad molecular expression profile in pluripotent stem cells, including many pathways targeted by clinical therapeutics, makes the use of pluripotent hiPSCs an attractive foundation for compound and drug screening.

In this report, we describe the establishment and characterisation of a laboratory-based drug screening platform that compares drug responses between multiple hiPSC lines in the pluripotent state. The platform combines a Cell Painting assay with advanced analytics and mass-spectrometry (MS)-based proteomic profiling to detect and explain variable drug responses. We validate the use of this screening platform using different clinically-approved drugs that target diverse physiological pathways and disease areas.

## RESULTS

### Establishing Pluripotency in hiPSC Cohorts in High-Throughput Format

To test whether we could detect and characterize variable drug responses across a cohort of hiPSC lines from different donors, the high throughput Cell Painting assay (*16*) was first adapted for use with hiPSC lines grown in the pluripotent state. We reasoned that an image-based Cell Painting assay could be used to detect variable responses and stratify cell lines accordingly, followed by using quantitative MS-based proteomics to identify potential mechanisms causing the variable responses (see Figure 1A).

**Figure 1.**
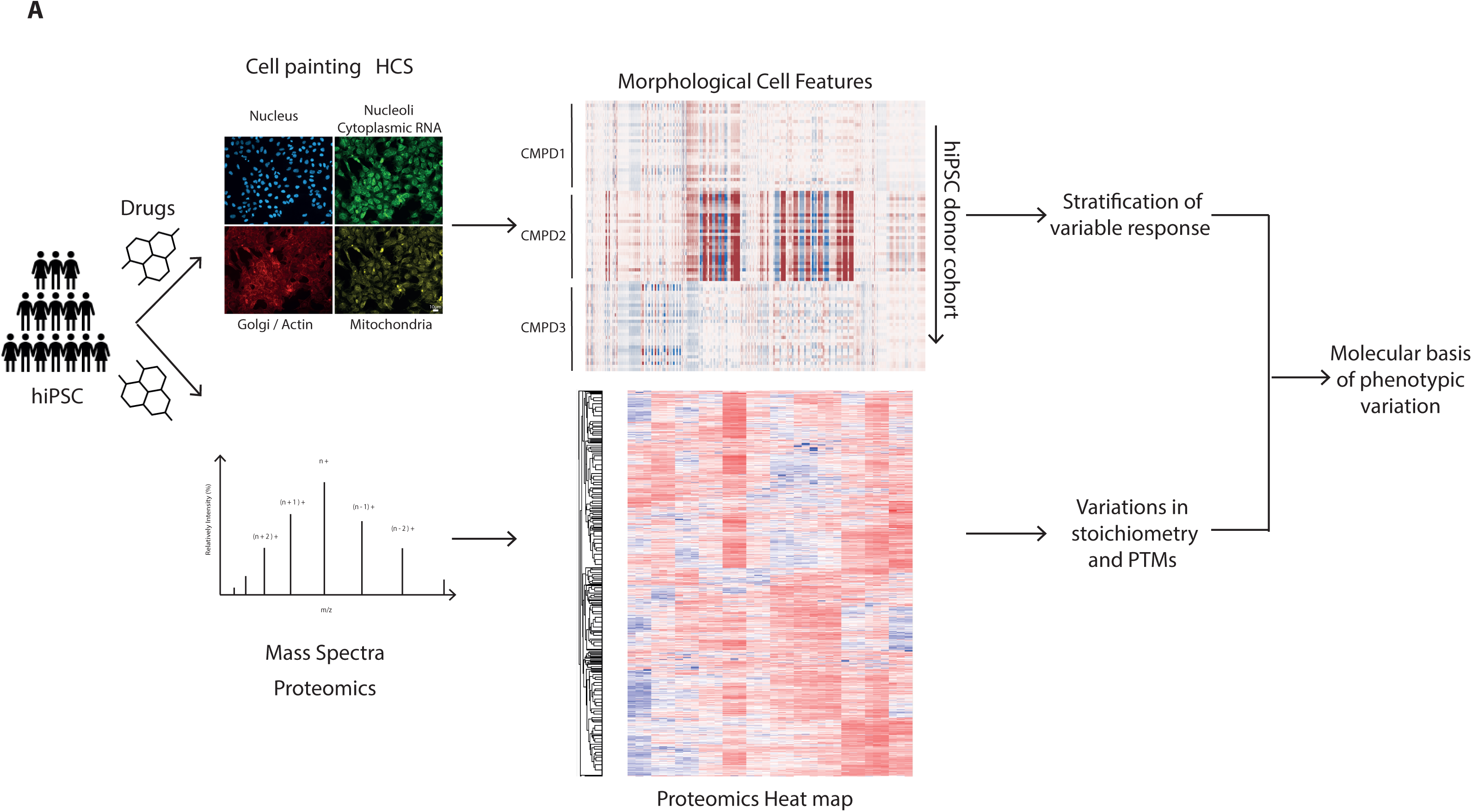
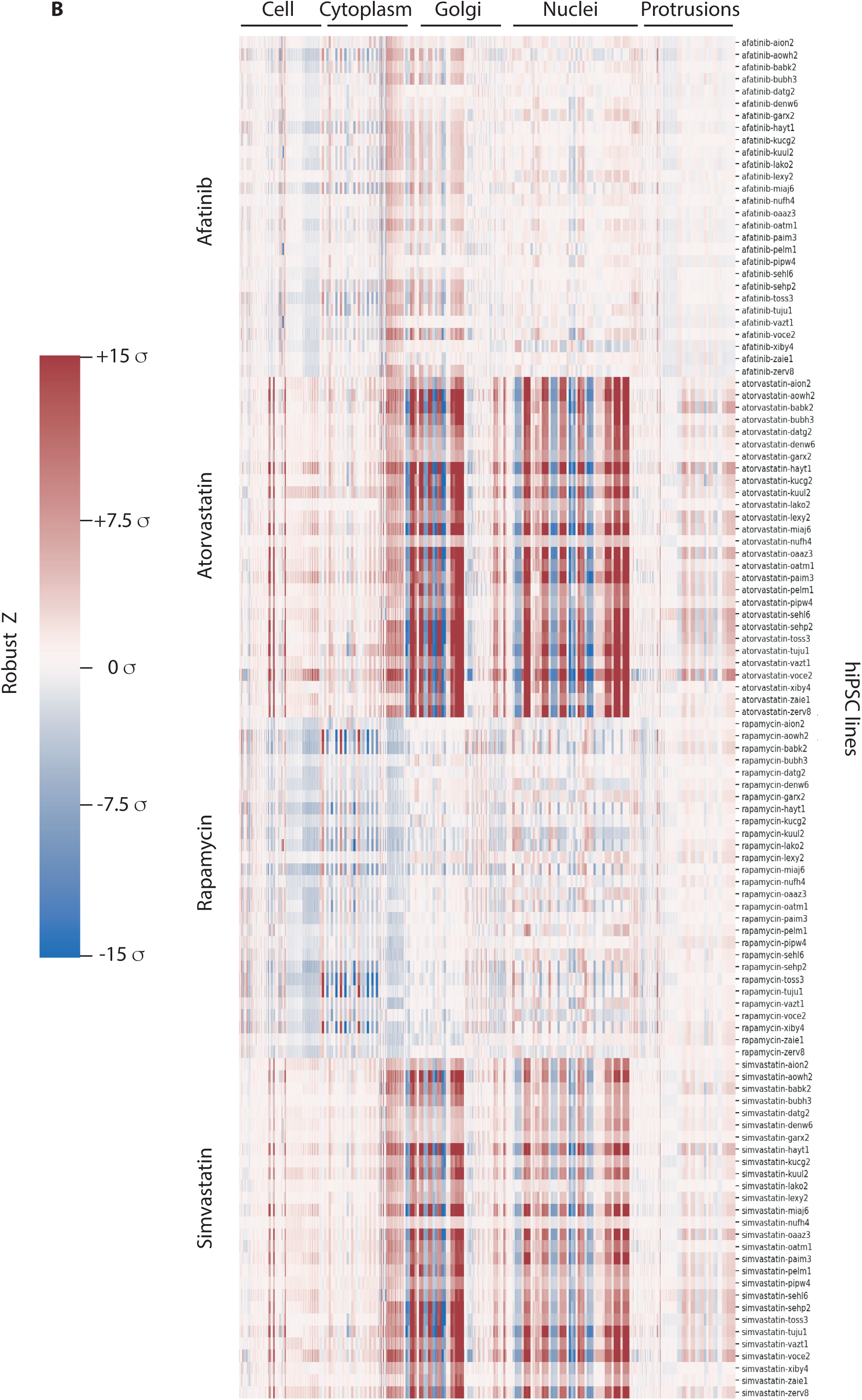
(A) Study overview. Quality controlled hiPSC lines from 43 donors in their pluripotent state were used for Cell Painting and proteomic analysis, by mass spectrometry, following addition of 52 FDA approved drugs. 38.2 million cells were imaged across all donors and 871 morphological traits were quantified per donor/drug combination. Variability of drug response of the hiPSC cohort was studied using both phenotypic screening and proteomics. (B) Heatmap of RobustZ scores from Cell Painting shows variations in the magnitude of drug responses across hiPSC lines. Differential responses to afatinib, atorvastatin, rapamycin and simvastatin are shown. The response to each drug creates a characteristic pattern of features, but with differences in the magnitude of the response. Color bar shows RobustZ scores normalized to DMSO expressed as standard deviations.

First, we established whether conditions could be established for culturing multiple hiPSC lines in the pluripotent state in 384 multi-well plates, suitable for high content imaging and analysis. Twenty-eight independent hiPSC lines, all derived from different donors (Supp Table 1, “Run1”), were propagated in mTeSR medium, plated at 3×10^4^ cells/cm^2^ in 384 multi-well plates for 72 hrs and then fixed and analysed by immunofluorescence to detect the established markers of pluripotency, i.e., Oct-4, Sox-2, Nanog and TRA-1-81 (Supp Figure 1A). All four markers showed strong, homogeneous staining.

We conclude, first, that multiple hiPSC lines can be cultured successfully in multi-well format suitable for large-scale, compound profiling by image analysis and second, that the panels of hiPSCs can be maintained in a pluripotent state throughout the 48-72 hours required for conducting the screening protocol.

### Using Cell Painting to Profile Variable Drug Responses

Next, the set of 28 independent hiPSC lines from different donors, (hereafter, “donors”), were grown in 384 multi-well plates for 24 hrs (see Methods for details), then treated either with compounds from a library of FDA-approved drugs, or with DMSO carrier, each at a single concentration (Supp Table 2). After incubation for a further 24 hrs in the presence of the drugs, the cells were fixed and stained with Cell Painting markers (see Materials and Methods). Images were recorded in an automated, plate-based imager, then imported into OMERO (9). Images were segmented with respect to the ‘nuclei’, ‘cytoplasm’, ‘Golgi’ and ‘protrusions’, using a custom CellProfiler pipeline (*17*). For each drug-donor combination, image features were calculated, using 871 separate feature measurements, from a total of 6,000-8,000 cells per donor-drug pair. After Robust Z’ normalisation (*18–20*), all measured features were scaled as the number of standard deviations different from DMSO for each of the hiPSC lines (*16*).

Figure 1B shows a heatmap of the measured features of the 28 donors, comparing treatment with 4 of the drugs from the library, i.e., atorvastatin, simvastatin, rapamycin and afatinib. The heatmap shows that each drug has characteristic feature patterns, but that different donors show varied magnitudes of response, visible as either darker, or lighter, horizontal stripes, respectively (Figure 1B). The feature patterns for atorvastatin and simvastatin are similar, consistent with them being competitive inhibitors of the same protein target, i.e., HMG CoA reductase, the rate-limiting enzyme of the cholesterol biosynthetic pathway. Interestingly, atorvastatin, which has a higher target affinity, also shows a stronger response than simvastatin in this assay.

These data demonstrate that the Cell Painting assay, which was originally developed for analysis of immortalised cancer lines (*16*), also generates characteristic phenotypic fingerprints in hiPSC lines. Moreover, the data show that Cell Painting fingerprints from hiPSC cohorts reveal variable responses between the lines from different donors. The major variation corresponds to differences in the magnitude of features.

Next, the assay was expanded to profile the hiPSC cohort responses to 52 FDA-approved drugs, which are used to treat a range of different diseases (Supp Table 2). A subset of the drugs tested are shown for visibility with full data available on Image Data Resource IDR (*21*). To remove highly correlated features, we calculated the Spearman correlation coefficient, *s*, and removed all features where *s* > 0.98, which left 442 filtered features for further analysis.

Figure 2A shows a heatmap of the Cell Painting induction, calculated as the fraction of filtered features with RobustZ > 2σ from DMSO (i.e. significant with >95% certainty) (*22*). This effectively reduces the filtered features for each drug-donor pair to a single statistic, which may underestimate the complexity of phenotypes. Despite this limitation, Figure 2A shows clearly that, first, hiPSC lines derived from different donors exhibit different levels of induction when treated with the same drugs and second, that examples of such differential response are observed between cell lines for all of the 52 different drugs tested.

**Figure 2.**
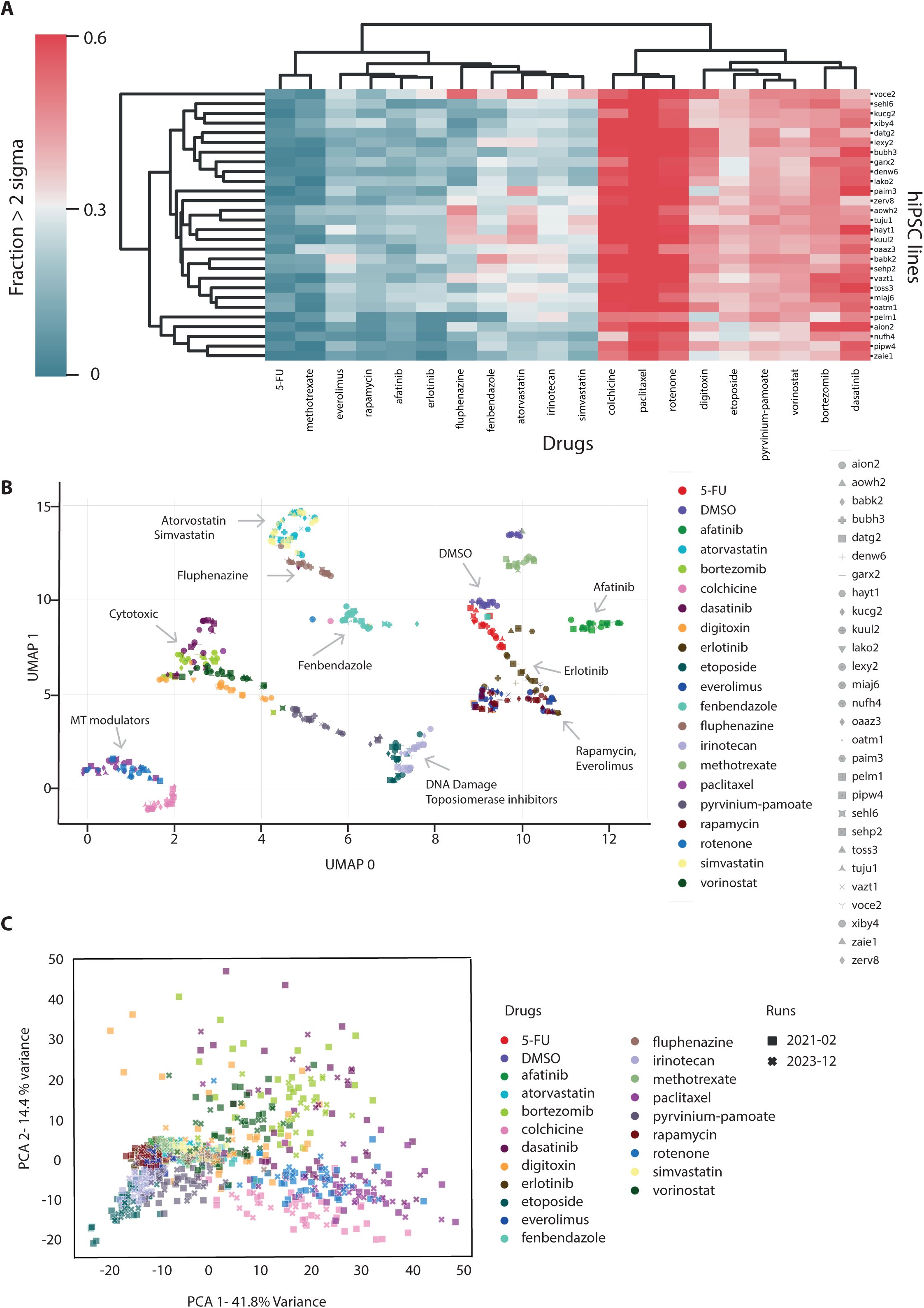
(A) Induction plot of drug responses for each hiPSC line shows variations in response magnitude. Induction is the fraction of features for each combination of drug and hiPSC line that differ in value by > 2σ (standard deviation) away from the DMSO control. (B) UMAP embedding of Cell Painting features measured for 28 hiPSC lines treated with different drugs. Cell lines are represented on the graph with different shapes and drugs with different colours. Representative separated clusters are indicated by arrows. For visibility, a subset of drugs used in this screen are shown. (C) PCA analysis of Cell Painting features measured for human iPSC lines from phenotypic screens performed in 2021-02 (28 lines) and 2023-12 (18 lines). Runs are represented on the graph with different shapes (“Run 2021-02” solid squares and “Run 2023-12” crosses) and drugs with different colours. The reproducibility of the assay results on different dates is indicated by the overlap of drug treatments in the different assay runs.

In all but one case, the individual cell lines show varied responses and appear to be either hyper- or hyposensitive to specific drugs. The exception was the cell line voce2, which showed a higher induction level to several, but not all cell lines. Combining the results shown in Figure 1B (feature heatmap) and Figure 2A (induction heatmap), we conclude that a significant source of variable responses between donor cell lines seen in the Cell Painting assay results from differences in the magnitude of response shown by each line.

We next asked if phenotypes caused by different compounds could be distinguished across the cohort of hiPSC lines. The filtered features from all drug-donor pairs were subjected to embedding with UMAP, which fits a manifold to a high-dimensional dataset and then projects this to a 2D plane (*23*) (Figure 2B). This showed that compound responses clustered according to their known mechanism of action. For example, statins, everolimus and rapamycin, topoisomerase inhibitors, cytotoxics and microtubule depolymerizers all formed separate, individual clusters. Interestingly, fluphenazine, an antipsychotic that acts by blocking specific dopamine receptors in the brain and that is used to treat schizophrenia and other psychotic disorders, clustered near the statins. While clustering in UMAP is not always a reliable indicator of similarity, we also noted the pattern of induction scores for fluphenazine was similar to that observed for atorvastatin (Figure. 2A). These results are consistent with reports that fluphenazine, like statins, affects lipid metabolism in human patients(*24*).

Besides showing that many drugs with similar modes of action cluster in this assay, the data also reveal cell lines that show variable responses to the same drugs. For example, when treated with rotenone, which disrupts microtubules (*25*), most hiPSC lines cluster together (“MT Modulators, Figure. 2B). An exception is the nufh4 cell line (pink and blue dot), which clusters instead with hiPSC lines treated with fenbendazole. This is a drug which also causes microtubule depolymerisation, but by a different mechanism to rotenone, which inhibits microtubule assembly dynamics (*26*) (filled circles with trefoil, Figure 2B). The nufh4 line consistently shows a lower induction score when treated with microtubule deploymerizers (Figure. 2A), which is consistent with nufh4 having a different response to rotenone than the other hiPSC lines.

To test whether these results were reproducible, a selection of the previously analysed hiPSC lines (Supp Table 1) were regrown from frozen stocks and analysed in Cell Painting assays after treatment with the same set of FDA-approved drugs, but this time using a different High Content Screening (HCS) imaging system (data collected on a Yokogawa CV7000 instead of InCell 2200). Using UMAP for visualisation, we observed similar clustering of Cell Painting features generated in this repeat set of assays to those collected from the original experiment that was performed 6 months previously, using independent cultures of each hiPSC line and with data acquired using a different type of HCS imaging system (Supp Figure 1B).

To further asses the reproducibility of the assay, we compared the measured phenotypes of hiPSC lines in an additional run of the assay, executed after an interval of 2 yrs 8 months (both datasets were collected on the same InCell 2200 imager) but with a different cohort of hiPSC lines (Supp Table 1). Figure 2C shows the comparison of the two assays, using Principal Component Analysis (PCA). This further validates the comparison between the two different runs and also avoids the effects of clustering in high-dimensional space that can affect UMAP (*27*). PCA analysis shows that similar compounds are indistinguishable in the two assay runs. Supplemental Figure 1 (C,E) shows a comparison of the two assays and again reveals that phenotypic responses are similar. As expected, PCA and UMAP show different clustering properties and different variations across cell lines, but the specific clustering responses of cell lines treated with different classes of drugs, as well as the variable responses observed to the same drug, between cell lines derived from different donors, are reproducible in this assay format. We conclude this represents intrinsic properties of the respective cell lines, independent of variables specific to individual assay runs, such as specific cell line stocks, prepared media, or the instrumentation used for image acquisition.

While many of the drug treatments resulted in tight clustering of the responses of different cell lines on the UMAP and PCA plots (Figure 2), a subset of the drugs tested, i.e. statins, 5-FU and erlotinib, showed more dispersed clustering of cell line responses. We hypothesised that this behaviour reflected an increased degree of variable response to these drugs between the respective cell lines. To test this hypothesis, we calculated the Euclidean distance of the feature vectors measured for each compound and DMSO, for each donor. Supplemental Fig 3B shows the result of this calculation, along with the mean and coefficient of variation (CoV) of the Euclidean distance for each drug across donors. This metric is elevated for simvastatin, atorvastatin, erlotinib and fenbendazole, consistent with the evidence from UMAP and the induction plot that response to these drugs varies across the donors (*27*).

The variable responses shown in Figure 2 and Supplemental Fig 3B might be due to intrinsic variations within an assay. To directly assess the variations in Cell Painting features in different assays, we calculated the Earth Mover Distance (EMD, see Methods) which has been shown previously to provide a useful measure of differences between phenotypic features in HCS assays (*28*). Supp Fig 2A shows the distribution of EMDs for all features from all compound treatments common between these two assay runs. Most show EMDs similar to DMSO, suggesting the differences are in the range of the assay controls. Cytotoxic compounds (e.g., bortezimib, dasatinib, colchicine) show a broader distribution of EMDs, which is likely due to cell death caused by these compounds during incubation. Inspired by the overall similarities in features between the two assays, we calculated induction scores for the drugs and donors that are common across these assays. Comparison of heatmaps of these induction scores shows the variations between drugs, but also between donors within individual drugs, are similar (Supp Fig 2B,C). These data suggest that the variations we observe in drug responses (e.g., Figures 1 and 2) across hiPSC lines generated from different donors are robust and reproducible.

### Linking Variable Responses to Protein Expression & Stoichiometry

Our previous studies on this HipSci cohort have demonstrated that variations in protein expression levels between lines derived from different donors can be controlled by specific genomic loci, i.e., protein Quantitative Trait Loci (pQTLs) (*13*). This suggested the hypothesis that a potential source of differential drug responses between hiPSC lines might be due to epistatic interactions, i.e., resulting from genomic variation between donors causing differences in protein expression that can modulate drug-induced phenotypes, for example by affecting the stoichiometry of components of drug response pathways (*29*). Importantly, this hypothesis takes into account drug-cell interactions outside of the direct interaction of a drug with its protein target. Furthermore, it can potentially explain why patterns of drug response features measured by Cell Painting appear similar across cell lines, while the magnitude of responses between lines differ, as seen in the feature and induction heatmaps (Figures 1B and 2A).

To test this hypothesis, we first used the induction plot (Figure 2A) and a graph of induction values (Supp Figure 3A), to classify cell lines as, respectively, “low” and “high” responders, for two well-characterised drugs, i.e., atorvastatin and simvastatin. These drugs share the same target, but are well-known to have different responses and efficacies in different individuals (*30*). Representative cell lines denw6 and zaie1 (“low responders”) and hayt1 and tuju1 (“high responders”), were each treated separately with either DMSO (control), or with either atorvastatin, or simvastatin, each at 5 µM, for 24 hrs, as in the Cell Painting assay. Following treatment, cell extracts were prepared and proteomes analysed by LC-MS/MS (see Methods). From the resulting data, differences in protein expression levels between DMSO- and drug-treated lines were visualised in ‘volcano plots’ (Figure 3). Combining data for treatment of all four cell lines (tuju1, hayt1, denw6, zaie1), with either atorvastatin (Figure 3A), or simvastatin (Figure 3B), showed that many of the proteins whose expression increased after drug treatment are involved in lipid metabolism (red dots in Figure 3A, B). Further, GO term enrichment analysis also showed a significant enrichment for expression of protein factors involved in lipid metabolism (Supp Table 3 and 4).

**Figure 3.**
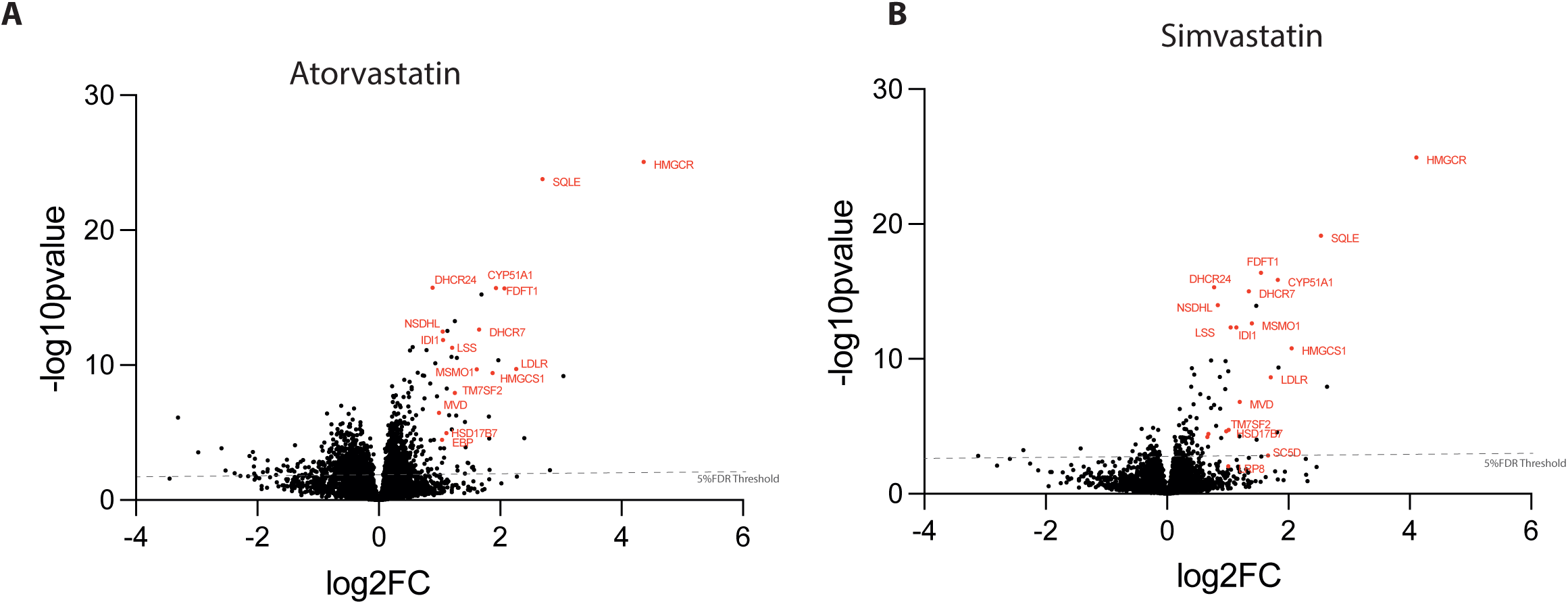
Volcano plots showing differential protein expression relative to DMSO in all four (tuju1, hayt1, denw6 and zaie1), hiPSC lines, treated with either atorvastatin (A) or simvastatin (B). The protein expression fold change (FC) and p value were calculated using Perseus. All sample runs (n=12), from each drug, were considered as one group, and the DMSO control set (n=12) was considered as a separate group. Representative proteins that play a role in the HMG-CoA reductase pathway are highlighted in red and show increased expression following drug treatment.

These data suggest that homeostatic mechanisms in the hiPSC lines respond to drug mediated inhibition of cholesterol metabolism by increasing the expression of enzymes required for cholesterol synthesis. While a common type of response to inhibition of cholesterol synthesis is seen across the hiPSC lines, differences were evident in the magnitude of responses between individual cell lines. This is illustrated by the response to either atorvastatin, or simvastatin, as detected by Cell Painting induction (Figure 2A). We therefore tested whether variation in response magnitude might be linked to differences in protein expression between the respective high and low response lines. Volcano plots comparing protein expression in the high and low response lines, after treatment with either atorvastatin (Figure 4A), or simvastatin (Supp Figure. 4A), show that multiple proteins involved in the cholesterol synthesis pathway (highlighted in purple), together with some other classes of proteins, have increased expression specifically in the high response lines. These data are consistent with the hypothesis that differential protein expression may explain, at least in part, the differences in the magnitude of drug responses between individual hiPSC lines.

**Figure 4.**
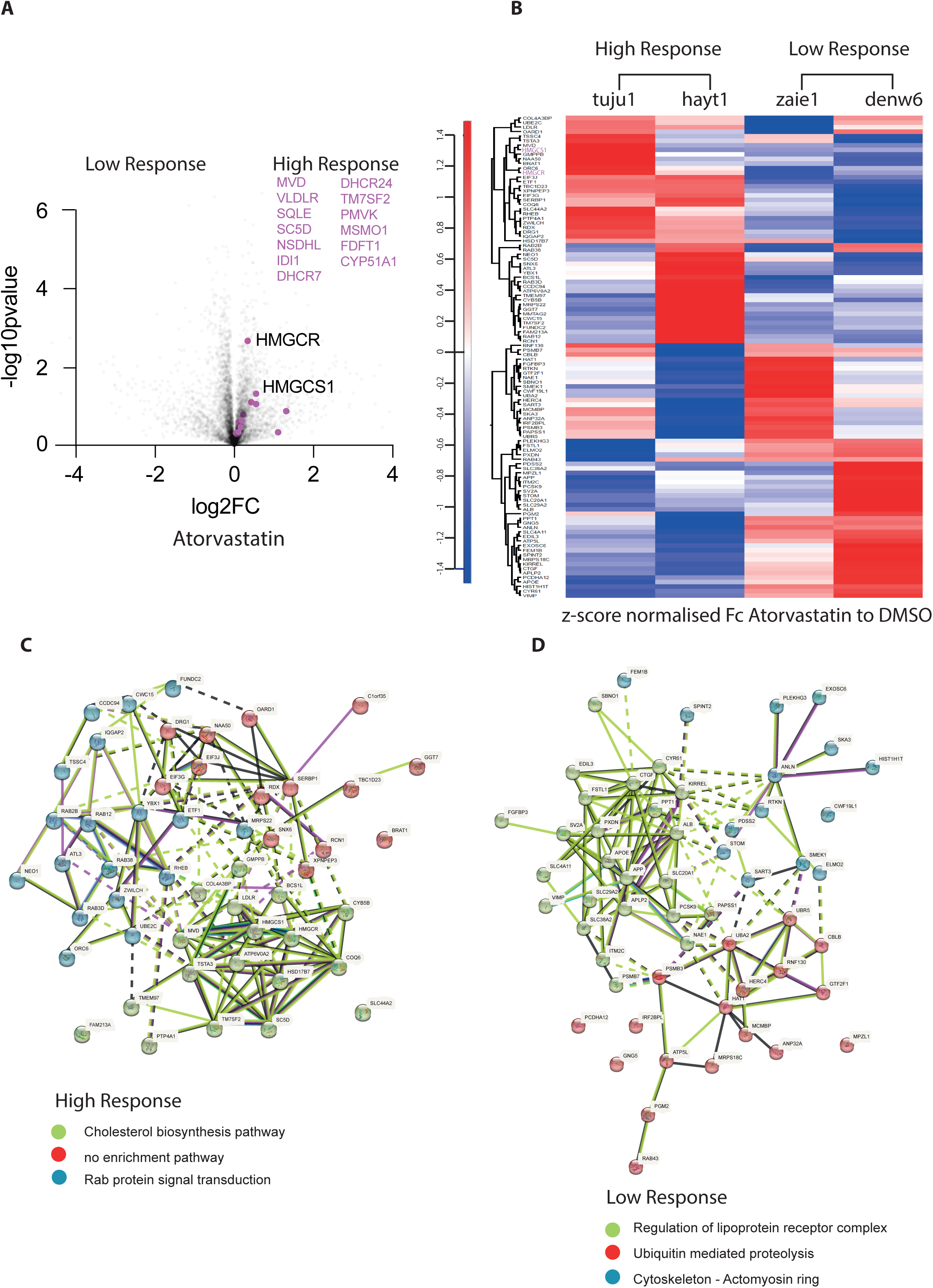
(A) Volcano plot of differential protein expression in four hiPSC lines, stratified by their respective high, or low, response to atorvastatin (see Supp Fig 3A). High response lines: tuju1 and hayt1. Low response lines: denw6 and zaie1. Protein FC and p value for each cell line were calculated as in Fig. 3. Sample runs from the drug treatment set (n=6), were considered as one group and the DMSO control set (n=6), were considered as a separate group. Members of the cholesterol biosynthesis pathway are represented as purple dots and listed on the right side of the graph. (A) Heat map analysis of proteins showing differential expression (log2FC_high-low_ >0.263) between the respective high and low response hiPSC lines, following atorvastatin treatment. Heatmap generation steps: the protein fold change (FC_high_) and p value (P_high_) for high response cell lines were calculated using sample runs from both high response cell lines to Atorvastatin set (n=6), considered as one group and the DMSO control set (n=6) considered as a separate group. The protein fold change (FC_low_) and p value (P_low_) for low response cell lines were also calculated. Only proteins with both P_high_ ≤0.05 and P_low_≤0.05 were considered further to compare the difference between high and low response. log2FC of these filtered proteins was calculated as (log2FC_high-low_= log2FC_high_-log2FC_low_). Heatmap shows FC between drug and DMSO control of the filtered list z-score normalised. StringDB analysis of filtered proteins shown in (B), in high response lines (C) and low response lines (D).

Next, we compared the expression of individual proteins in each of the four drug-treated lines (Figure 4B). Proteins were identified whose expression differed significantly in high vs low responders (|log_2_FC| >0.263 and *p* < 0.05, see Materials and Methods). This reveals clear differences in the responses to the two statin drugs, suggesting that there may be variation in the mechanisms of response to statin treatment between each hiPSC line (see also Supp Figure 4B, Supp Table 9).

Interestingly, changes in protein expression in the high response lines include increased expression of HMGCR and HMGCS1, which are directly involved in the cholesterol mevalonate synthesis pathway. In contrast, both “low response lines”, show increased expression of proteins involved in regulating the stability and recycling of lipoprotein receptor complexes. This includes PCSK9, a protein that provides post translational regulation of LDL receptor and directs it for lysosomal degradation, rather than regular recycling potentially limiting the efficacy of statins. In addition, SCARB1, a cell surface HDL receptor that mediates HDL-cholesteryl ester uptake, as well as ApoE, a plasma protein that participates in the transport of cholesterol and other lipids between cells and has been linked to the effect of statins (*31–39*). Lipoprotein receptor complexes play an important role in the regulation of plasma and intracellular cholesterol homeostasis(*40*). These differences in protein expression between the respective high and low responder hiPSC lines to statin treatment suggest that variable activity of biosynthetic and/or influx mechanisms may be involved in determining how each hiPSC line responds to statins. In addition to identifying well-established targets of statins, characterising the proteins whose expression increases in the high response lines, using GO enrichment and STRING analysis (https://string-db.org/), identified factors involved in Rab protein signal transduction and in the metabolism of RNA (Figure 4C and Supp Figure 4C; Supp Table 5 and 7). These are all known to change expression upon statin treatment (*41–45*). In contrast, the low response lines showed increased expression of proteins linked to the cytoskeleton, along with several ubiquitin E2 and E3 enzymes, suggesting that ubiquitin-conjugation levels may affect the degree of statin response (Figure 4D and Supp Figure 4D; Supp Table 6 and 8).

These data support the view that a diversity of factors and pathways can affect the cellular response to well-defined drug perturbations and these differences are due, at least in part, to population-level variations in gene and protein expression between the respective lines.

To explore further the observed variable drug responses, we calculated the pairwise correlation matrix for the induction metric (see Figure 2A) for 32 drugs across donors. This provides a measure of the pattern of variable response across a given set of donors. Supp Figure 5 shows a visualisation of these data in a heatmap and identifies sets of compounds that share common patterns of variable response. For example, erlotinib and afatinib, both ErbB family receptor inhibitors and the statins and fluphenazine, behave similarly. Everolimus, etoposide and irinotecan share a similar variable response pattern, which may be linked to a role for the mTor pathway in controlling the DNA damage response (*46, 47*). In addition, warfarin, omeprazole, niclosamide and metoclopramide, which have very different targets, nonetheless share a similar variable response pattern across hiPSC lines. We hypothesise that this may be linked to a shared off-target effect, e.g., mitotoxicity (*48*). This is an interesting point to explore in further studies in the future.

## DISCUSSION

We have established, to our knowledge, the first laboratory pipelines for the systematic analysis of population-level variable drug responses, using a cohort of hiPSC lines, derived from multiple, healthy donors. Previous work has described culturing hiPSCs from a single patient in 384-well plate for drug screening, In this study, we have used a cohort of hiPSC lines from many different donors and Cell Painting to garner significantly more information from the screens (*12, 49*). This pipeline provides a ‘genetically-informed’, high-throughput assay for identifying and characterising variable drug responses in human cells, using Cell Painting and quantitative proteomics. This analysis provides an initial quantitative measure of variable response across a cohort of hiPSC lines.(*50*)

The choice of screening hiPSC lines specifically in their undifferentiated, pluripotent state leverages the recent discovery that, at least in part, gene expression in hiPSC lines reflects the genetic identity of the donor at both the RNA and protein levels (*12, 13*). Thus, the undifferentiated hiPSC lines can behave as avatars of the donors, with respect to their proteomes and cellular phenotypes. The corollary is that drug screening assays, using cohorts of pluripotent hiPSC lines from different donors, allows objective stratification based on measurements of phenotypic responses. We hypothesise that these data may reveal important information about variable human drug responses that is germane to clinically relevant differential drug responses between patients. It is important to note that the HipSci cohort covers limited ethnic diversity, with a high concentration of ethnic European donors (*51*). In the future, we aim to expand our working cohort to include hiPSC lines from a broader range of donor ethnicity, which may provide an opportunity to detect variable efficacy and toxicity of drugs in different ethnic groups (*50, 51*).

The data presented in this study show that features can be extracted from the analysis of high-throughput, Cell Painting, fluorescence microscopy images and used to stratify differences in the magnitude of drug responses between different hiSPC lines, (illustrated in Figures 1 and 2). The data also show that the stratification of cell lines, based upon their degree of drug response, is reproducible in this assay format. Further, by using quantitative proteomic analysis to compare the stratified sets of hiPSC lines we show that additional information relevant to understanding the mechanisms underlying variable drug response phenotypes can be identified. We have focussed on proteome level analysis here because proteins are the direct targets of most drugs in clinical use and because proteins are also the primary mediators of most disease processes and mechanisms of drug action.

We propose a hypothesis to explain the reproducible, differential response to drug treatment we have detected. This hypothesis postulates that variations between the cell lines, at the genomic and/or epigenomic level, in turn determine the respective proteomes and thereby lead to differences in drug-cell interactions. We envisage that drug-cell response phenotypes can reflect multiple epistatic interactions that are predominantly mediated at the proteome level. This hypothesis is supported by our comparison of proteomes of hiPSC lines that were stratified by Cell Painting as either ‘high’, or ‘low’ responders, respectively, to treatment with either atorvastatin and simvastatin. While we have concentrated here on comparing protein expression levels between cell lines, we note that other comparative analyses, for example comparing protein phosphorylation and other protein post translational modifications (PTMs) and protein-protein interactions, can also provide valuable insights into the mechanisms causing variable drug responses between the different cell lines.

In this study, we have investigated potential mechanisms involved in the variable responses seen to statins in this screening assay. Statins are drugs developed to reduce the level of cholesterol in blood and thus lower the risk of developing cardiac and circulatory diseases, including angina, heart attack and stroke (*52*). Separate, but interconnected pathways, control cholesterol homeostasis, including cholesterol influx, mediated by lipoprotein receptors, cholesterol synthesis, regulated by HMG-CoA reductase and the mevalonate pathway and cholesterol efflux, mediated by transport proteins. Each of these pathways respond to multiple inputs that collectively define cholesterol levels (*53*). In addition, our data suggest that modulation of protein stability by Ub-mediated proteolysis is also an important factor in cholesterol homeostasis (Figure 4D). Even within the relatively limited sample of different donor lines we have analysed so far, we see evidence of varied regulatory effects. The comparative proteomic analysis supports the view that “high response” lines can be determined by modulation of the endogenous cholesterol synthesis pathway, while “low response lines” may instead reflect primarily regulation of the cholesterol influx and efflux pathways.

In conclusion, we have established and validated a flexible, laboratory-based high-throughput screening platform that can detect and characterise examples of variable drug response for clinically relevant therapeutics. We anticipate that the use of this platform will, in the future, help to improve the success rate of clinical trials by predicting variable responses and providing evidence of biomarkers that may improve the selection of suitable patients most likely to benefit from new drug candidates. We also anticipate that the use of carefully selected hiPSC cohorts, derived from donors of different sex and genetic backgrounds, can help to improve the development of safe new drugs that are suitable for the needs of modern, diverse human societies.

## Acknowledgements

We thank members of the National Phenotypic Screening Centre and Human Pluripotent Stem Cell Facility at the University of Dundee for invaluable help and advice. We thank University of Dundee’s School of Life Sciences for their support of this project.

## MATERIALS & METHODS

### Cell Culture

Human iPSC lines used in this study were from the HipSci cohort (4). Feeder-free human iPSC lines were cultured in Essential 8 (E8) medium (E8 complete medium supplemented with (50x) E8 supplement ThermoFisher-A1517001) on tissue-culture dishes coated with 10 µg/cm^2^ of reduced Growth Factor Basement Membrane matrix (Geltrex, ThermoFisher A1413202 resuspended in basal medium DMEM/F12 Thermo Fisher 21331020). Medium was changed daily.

To passage feeder-free hiPSC lines, cells were washed with PBS and incubated briefly (3-5 min), with 0.5 mM PBS-EDTA solution. After PBS-EDTA solution was removed, cell clusters were resuspended in E8 medium and seeded at ratios of 1:4 to 1:6, depending on their confluency, on Geltrex-coated tissue culture dishes. Once established, undifferentiated, feeder-free human iPSC lines were expanded and aliquots frozen for future use, prior to transition to TeSR medium for further study. To transition to TeSR medium supplemented with bFGF (Peprotech 100-1813, at a final concentration of 30 ng/ml), Noggin (Peprotech 120-10C, at a final concentration of 10ng/ml), Activin A (Peprotech 120-14P, at a final concentration of 10ng/ml), E8 medium of hiPSC lines was exchanged daily with medium containing increasing amounts of TeSR medium, until human iPSC lines were completely maintained in TeSR medium alone, typically over 1-2 weeks.

To passage human iPSC lines after transition to TeSR medium, cells were washed with PBS and incubated briefly with TrypLE Select (ThermoFisher 12563029), to create a single cell suspension prior to resuspension in TeSR medium supplemented with ROCK kinase inhibitor (Tocris 1254, at a final concentration of 10 µM), to enhance single cell survival, bFGF (Peprotech 100-1813, at a final concentration of 30 ng/ml), Noggin (Peprotech 120-10C, at a final concentration of 10ng/ml), Activin A (Peprotech 120-14P, at a final concentration of 10ng/ml) and seeded at a concentration of 5×10^4^ cells/cm^2^ on Geltrex-coated tissue culture dishes. 24 hrs after replating Rock kinase inhibitor was removed from the culture medium. Pluripotency of TeSR-transitioned hiPSC lines was verified by immunofluorescence (see below) prior to either cell banking, or experimental use. Cells were grown at 37°C and 5% CO_2_. Medium was changed daily.

hiPSC line ID number and details of the cell lines are listed in Supp Table 1.

The data from the experimental runs are included in this report. Run 2021-02, 52 drugs, 28 donors 1,456 drug-donors pairs). Run 2021-10, 8 donors, 52 drugs (416 drug-donor pairs). Run 2023-12, 18 donors, 52drug (936 drug-donor pairs).

### Cell Staining

384-well tissue culture plates (CellCarrier-384 Ultra Microplates, Perkin Elmer 6057300) were coated with human plasma fibronectin (Merck Millipore FC010) at a concentration of 5 µg/cm^2^. Cells were passaged as described above with TrypLE Select, counted and seeded on the fibronectin-coated wells, at a concentration of 3×10^4^ /cm^2^. Cell line plating was in rows, with six wells per condition for each cell line (six technical replicates). Each replicate was sampled at 9 locations and repeated in a separate biological replicate separated by a week (i.e. 2 bioreplicates per assay run).

Cells were incubated for 24 hrs prior to drug treatment, followed by a further 24 hrs incubation before final fixation, staining and high content imaging. Antibody staining of a separate 384-well plate, at 48 hrs, confirmed that pluripotency was maintained at 48 hrs.

Mitotracker (Thermo Fisher M22426), staining was performed in live cells with a 30 min incubation at 37°C prior to fixation. Fixation, permeabilisation and immunostaining were performed at room temperature. 8% Paraformaldehyde in PBS (PFA, Sigma F8775), was added to an equal volume of cell culture medium in wells of a 384-well plate for a final concentration of 4% Paraformaldehyde and incubated for 15 min. Fixed cells were washed in PBS and permeabilised with 0.1% v/v Triton-X100 (Sigma) for 15 min. Following permeabilisation, cells were washed in TBS and stained with DAPI (Thermo Fisher D1306 at 5 µg/ml), SYTO 14 (Thermo Fisher S7576 at 2.5 µM), Concanavalin A (Thermo Fisher C11252 at 20 µg/ml), Phalloidin Alexa (Thermo Fisher A12381 at 66 nM), and WGA (Thermo Fisher W11262 at 11.8 µg/ml) for 1hr. Staining volume was 30 µl/well. Cell fixation, permeabilisation, staining and washing steps (2x, wash volume 50 µl) were performed using automated liquid handling systems (405 plate washer, BioTek– Tempest, Formulatrix).

### Drug treatment

Drug dosing on 384-well plates was performed using an Echo acoustic liquid handler and corresponding software (Labcyte). Screened compounds were selected from the appropriate chemical library plates containing either Cloud library (Enamine), or an in-house library of publicly available compounds. Drug dosing was performed at final concentrations of; etoposide 1.7 µM, Berberine Chloride 2.68 µM and Fenbendazole 3.34 µM. For all other compounds drug dosing was performed at a final concentration of 5 µM. Supp Table 2 lists the compounds and their final concentrations used in this study. For proteomics analysis drug treatments with either simvastatin (Tocris 1965), or atorvastatin; (Selleckchem S5715) were performed in triplicate in 6-well plates at a final concentration of 5 µM for 24hrs.

### Cell staining and analysis of pluripotency markers

All cell lines used in this study were quality controlled for pluripotency prior and during the HCS assays. For each experiment 8 wells of each cell line were individually stained with pluripotency marker antibodies.

All fixation, permeabilisation and immunostaining analyses were performed at room temperature, apart from primary antibody incubation, which was carried out at 4°C overnight. 8% Paraformaldehyde in PBS (PFA, Sigma F8775), was added to an equal volume of cell culture medium in wells of a 384-well plate for a final concentration of 4% Paraformaldehyde and incubated for 15 min. Fixed cells were washed twice in PBS and permeabilised with 0.5% v/v Triton-X100 (Sigma) for 10 min (apart from TRA1-81 staining). Following permeabilization cells were washed twice in PBS, then blocked in 10% normal donkey serum for 1 hr prior to antibody incubations. Pluripotency marker antibodies used in this study are: Nanog, Oct4, Sox2, Tra-1-81 all at 1:500, (Oct4 (C30A3), Sox2 (D6D9), Nanog (D73G4) and TRA181 (TRA-1-81) Cell Signalling StemLight Pluripotency Antibody Kit 9656). All affinity purified donkey secondary antibodies (either Alexa 488 or Alexa 594) were purchased from Jackson Immunoresearch Europe. Hoechst 33342 (Thermo Fisher) was used to stain DNA. Images of cells (for each hiPSC line, 9 fields/well, 16 wells per pluripotency marker), stained with pluripotency marker antibodies were segmented based on DAPI signal and the integrated nuclear intensity of each marker was quantified using CellProfiler. Distributions of integrated intensities of all nuclear pluripotency markers were monotonic and correlated with integrated DAPI staining.

### Imaging

Imaging data were collected on an InCell 2200 high content imager, with a 20x/NA 0.45 Plan Fluor lens and a Quad2 multichroic mirror (GE, Issaquah, WA) or a CV7000 high content imager, with a 20x/0.45 lens at epifluorescence mode (Yokogawa). For each assay, we performed 6 technical replicates (each in separate wells), with each replicate sampled at 9 locations, and repeated in a separate biological replicate separated by a week (i.e 2 bio replicates per assay run). All of these data are then combined into a single well measurement, producing an n-dimensional data vector per well.

### Data Processing, Analysis and Visualization of Multiple Cell Lines

Raw images were imported into OMERO Plus (Glencoe Software, Inc., (*54*)) and then processed using a custom pipeline in CellProfiler (*16*), which segmented nuclei, cytoplasm, Golgi and cortical protrusions and calculated a range of defined features. All further steps were executed using the Pandas Python library (*55*). Features for each plate were Robust-Z normalised by the median of features for the DMSO control for that plate (*16, 56*).The Robust Z-score converts the population of features derived from individual cells to a distribution of zero mean and unit variance. Robust Z is calculated according to the formula x’ _i_ =(x_i_-µ)/μ where x _i_ is the measured value for each feature, x’ _i_ is the normalised output value and µ and μ are the median and median absolute deviation (*18–20*). Features with a coefficient of variation >0.50 were removed from further analysis. We further removed all features with |Spearman coefficient| > 0.98. Data were then visualised using Uniform Manifold Approximation and Projection (UMAP (*23*)). UMAP parameters were n_neighbours = 15, mindist = 0.01 and a cosine metric was used for scaling the distance between points. GO enrichment analysis was carried out using WebGestalt (WEB-based Gene SeT AnaLysis Toolkit) (*57, 58*). Functional protein-protein interaction analysis was carried out using STRING-DB (*59*).

Nearest neighbour distances were calculated using Euclidean distance in scikit-learn (*60*). The Earth Mover Distance (EMD) was calculated using sci-py (*61*) for filtered features from individual wells for each drug-donor pair across a pair of assays. The frequency distribution of EMDs for filtered features was calculated and plotted using Seaborn (*62*).

### Proteomics sample preparation

To support parallel MS-based proteomics analysis in this screening platform in a 6-well plate format, we also tested whether the expression of pluripotency markers was maintained when hiPSC lines were plated at 5×10^4^ cells/cm^2^, prior to any drug treatment and cultured as previously described. This showed that strong, homogeneous expression of the four pluripotency markers was maintained at this higher cell density (not shown).

All hiPSCs samples were washed twice with 1x PBS, on ice, then centrifuged at 300g to collect the cell pellets. 4% SDS lysis buffer (10 mM TCEP, 100 mM Tris-HCl, pH 7.4) was used to lyse the cells and extract proteins, with 1x Protease and Phosphatase Inhibitor (A32961, Thermo Fisher) added. 1 mL lysis buffer was added to the cell pellets. After 10 cycles of sonication, (30s on/off), using the Bioruptor® Pico, the protein samples were centrifuged at 20,000*xg* for 15 min. The supernatant proteins were denatured at 95°C for 10 min, then alkylated with 40 mM IAA (Iodoacetamide), (final concentration) and incubated in the dark at room temperature for 30 min. The protein concentration was measured using the EZQ™ Protein Quantitation Kit following the manufacturer’s instructions.

500 ug proteins from each sample were further processed using the SP3 protocol, as described (*63*). In brief, protein samples were mixed with SP3 beads (1:10, protein:beads) and digested with LysC/trypsin mixture (1:50, enzyme:protein). Peptides were eluted from the SP3 beads with 2% DMSO. The peptide concentration was measured using the Pierce™ Quantitative Fluorometric Peptide Assay following the manufacturer’s instructions. Peptide samples were stored at −20℃ before LC-MS/MS analysis.

### LC-MS/MS analysis

For each cell line, three biological replicates per condition were generated. They were combined in one sample, which was analysed in three separate, replicate MS runs. Each hiPSC line was treated three times with either vehicle control, or specified drug.

All peptide samples were analysed by using an Orbitrap Fusion Tribrid mass spectrometer (Thermo Fisher Scientific), equipped with a Dionex UltiMate™ 3000 RSLCnano System, Thermo Fisher Scientific). Peptides from each sample were loaded onto a 75 μm × 2 cm PepMap-C18 pre-column and resolved on a 75 μm × 50 cm PepMap-C18 EASY-Spray temperature-controlled integrated column-emitter (Thermo Fisher Scientific), using either a 160 min or a 280 min multistep gradient from 5% B to 95% B with a constant flow rate of 300 nl/min. The mobile phases are: H_2_O incorporating 0.1% FA (solvent A) and 80% ACN in H_2_O incorporating 0.1% FA (solvent B). The details for 160 min gradient are: 0-6 min, 1%-5% B; 6-145 min, 5%-38% B; 145-155 min, 38%-95% B; 155-160 min, 95%B. The details for 280 min gradient are: 0-6 min, 1%-5% B; 6-255 min, 5%-38% B; 255-270 min, 38%-95% B; 270-280 min, 95% B.

For total proteome analysis, the data were acquired under the control of Xcalibur software in data-independent acquisition mode (DIA). For each sample, one run of 280 min method with 97 DIA windows was performed for boosting the identification of proteins (*64*) and another three replicate runs of 160 min method were performed for the quantification of proteins.

For the 160 min gradient method, the survey scan was acquired in the Orbitrap covering the m/z range from 375 to 1,175 Da with a mass resolution of 60,000 and an automatic gain control (AGC) target of 4.0 × 10^5^ ions. MS/MS was acquired in the Orbitrap covering the m/z range from 200 to 2,000 Da with a mass resolution of 30,000 and an AGC target of 1.0 × 10^5^ and 60 ms maximum injection time. The sequential DIA isolation windows are 10 m/z from 375 to 775 and 40 m/z from 775-1175. Fragmentation was performed using a HCD collision energy of 30%. For the 280 min gradient method, the survey scan was acquired in the Orbitrap covering the m/z range from 355 to 1,150 Da with a mass resolution of 120,000 and an automatic gain control (AGC) target of 4.0 × 10^5^ ions. MS/MS was acquired in the Orbitrap covering the m/z range from 200 to 2,000 Da with a mass resolution of 30,000 and an AGC target of 1.0 × 10^5^ and 60 ms maximum injection time. The sequential DIA isolation windows are 15 m/z from 350 to 410, 10 m/z from 410 to 440, 15 m/z from 440 to 645, 5 m/z from 645 to 775, 10 m/z from 775 to 875, 15 m/z from 875 to 965 and 27 m/z from 965-1150. The fragmentation was performed using a HCD collision energy of 30%.

### DIA MS data analysis

For DIA analysis, the raw MS data of different sample runs from both drug and DMSO set were analysed together using DIA-NN v1.8.1(*65*). The in silico spectral library was generated by DIA-NN, using the Homo sapiens database from UniProt (SwissProt October 2021). The data were searched with the following parameters: stable modification of carbamidomethyl (C), N-term M excision, with a maximum of 1 missed tryptic cleavage threshold, match between runs (MBR) option checked. DIA-NN output reports were further processed with an R package, as described previously (*66*) to perform the MaxLFQ-based protein quantification, with precursors x samples matrix filtered at 1% precursor and protein group FDR. Statistical analyses were performed using Perseus v1.6.15.0 (*67, 68*) In brief, the protein quantification result was imported to Perseus and the copy number of each protein was calculated based on its intensity using a “Proteomic Ruler” (*69*). Only proteins with valid quantification values in at least 1/3 of total sample runs were considered for further comparison. Missing values were replaced from a normal distribution. After median normalisation, two sample student’s T tests were performed for p-value generation, using permutation-based FDR for truncation. Heatmap and K-means clustering analysis was performed based on the Euclidean distance, using the default settings in Perseus. In total, 5,895 proteins were quantified using DIA analysis and used for further analysis.

For heat map generation of Figure 4B and Supp Figure 4B: At the first step of differential expression analysis, a two sample-test using Student’s T-test with the default setting in Perseus was performed to determine if the means of two groups (drug and DMSO, for either high or low response cell lines) are significantly different from each other.

The protein fold change (FC_high or low_) and p value (P_high or low_) were generated, respectively. Proteins with p<0.05 were considered further and used to create the results show in Figure 4B and Supp Figure 4B. Z-score normalisation was used for Heat map analysis, according to the protocol in (*68*).

## Data availability

The mass spectrometry proteomics data have been deposited to the ProteomeXchange Consortium, via the PRIDE (*70*) partner repository with the dataset identifier PXD050194. Cell painting data will be deposited in the Image Data Resource (IDR).

## Code availability

All data analysis code used methods published by others (*16*) and tools from pandas (*71*), numpy (*72*), scikit-learn (*73*), SciPy (*61*).

