## Supplemental Info for "Screening for variable drug responses using human iPSC cohorts"

Platani et al

Supp Figure 1

(A) Maintenance of pluripotency in hiPSC lines in HCS assay format. Representative micrographs from six hiPSC lines, plated in 384-well format and stained with pluripotency markers (Oct4, Sox2, Nanog and TRA-181). Scale bar; 100µm.

(B) UMAP Embedding of Cell Painting features measured for eight human iPSC lines from phenotypic screens performed on two different dates (2021-02 and 2021-10 on two different imaging systems. hiPSC lines were treated with drugs as described (see Materials and Methods). Cell lines are represented on the graph with different shapes and drugs with different colours. Arrows point to representative drug clusters. Note the high degree of reproducibility of the assay results generated on different dates.

(C) PCA analysis of Cell Painting features measured for hiPSC lines in the 2021-02 assay run. Drugs are represented with different colours.

(D) PCA analysis of Cell Painting features measured for hiPSC lines in the 2021-10 run. Drugs are represented with different colours.

(E) PCA analysis of Cell Painting features measured for hiPSC lines in the 2023-12 run. (Run “2023-12” -18 hiPSC lines). Drugs are represented with different colours.

Supp Figure 2

(A) Plot of distributions of Wasserstein distances between each feature calculated from Cell Painting from the two assays shown in Supp Fig 1B-D. The data for these assays were collected in February and October 2021 respectively, on two different imaging systems (InCell 2200 and CV700 high content imagers). The Wasserstein distance provides a metric of similarity of two distribution (*1*). Wasserstein distances of features for each drug-donor pair were calculated and the frequency distribution of the medians of the features for each drug was then plotted. Different drugs show different distributions, with cytotoxic drugs (e.g., bortezimib, dasatinib, paclitaxel and colchicine) showing higher Wasserstein distances i.e., great variation between features in the two different runs.

(B) and (C) Induction plots of drugs and donors that were common across the two assays show in Supp Fig 1B-D. Values in the heatmap are calculated induction values using a cutoff of 2 sigma (same is in Fig 2A).

Supp Figure 3

(A) Bar graph of induction values for hiPSC lines treated with atorvastatin (orange) or with simvastatin (blue). Cell lines used for proteomic analysis are shown in grey boxes.

(B) Table of nearest neighbour analysis. Spearman correlation filtered features for all drugs were used to calculate Euclidean distance from DMSO. Induction mean, standard deviation and coefficient of variation (CoV) for each drug are shown.

Supp Figure 4

(A) Volcano plot of differential protein expression in four hiPSC lines, stratified by their respective high, or low, response to simvastatin (see Supp Fig 3A). High response lines: tuju1 and hayt1. Low response lines: denw6 and zaie1. Protein FC and p value for each cell line were calculated as in Fig. 3. Sample runs from drug treatment set (n=6) were considered as one group and the DMSO control set (n=6) was considered as a separate group. Members of cholesterol biosynthesis pathway are represented as purple dots and listed on the right side of the graph.

(B) Heat map analysis of proteins showing differential expression ($log2FC_{high-low} > 0.263$) between high and low response hiPSC lines following simvastatin treatment. Heatmap generation steps: The protein fold change ($FC_{high}$) and p value ($P_{high}$) for high response cell lines were calculated with sample runs from both high response cell lines to Atorvastatin set (n=6) considered as one group and the DMSO control set (n=6) considered as a separate group. The protein fold change ($FC_{low}$) and p value ($P_{low}$) for low response cell lines were also calculated. Only proteins with both $P_{high} \leq 0.05$ and $P_{low} \leq 0.05$ were considered further, to compare the difference between high and low response. log2FC of these filtered proteins was calculated as ($log2FC_{high-low} = log2FC_{high} - log2FC_{low}$). Heatmap shows FC between drug and DMSO of filtered list z-score normalised. StringDB analysis of filtered proteins shown in (B) in high response lines (C) and low response lines (D).

Supp Figure 5

(A) Pairwise correlation matrix of induction values (see Fig. 2A) for all drugs and across all (28) donors. Note the clusters formed based on the patterns of induction across the donor cohort, suggesting similar responses between different drugs.

### **Supplemental Tables**

Supplemental Table 1: List of hiPSC lines used in this study

Supplemental Table 2: List of drugs, disease targets and concentrations used in this study

Supplemental Table 3: GO enrichment analysis of top 5% proteins showing increased expression in all four hiPSC lines used for proteomic analysis following atorvastatin treatment. Note pathways related to lipid metabolism.

Supplemental Table 4: GO enrichment analysis of top 5% proteins showing increased expression in all four hiPSC lines used for proteomic analysis following simvastatin treatment.

Supplemental Table 5: GO Enrichment Analysis of gene sets/pathways and their associated p-values for highly expressed proteins (Figure 4 heat map (B) values >0.2) of high response lines following atorvastatin treatment.

Supplemental Table 6: GO Enrichment Analysis of gene sets/pathways and their associated p-values for highly expressed proteins (Figure 4 heat map (B) values >0.2) of low response lines following atorvastatin treatment.

Supplemental Table 7: GO Enrichment Analysis of gene sets/pathways and their associated p-values for highly expressed proteins (Supp Figure 4 heat map (B) values >0.2) of high response lines following simvastatin treatment.

Supplemental Table 8: GO Enrichment Analysis of gene sets/pathways and their associated p-values for highly expressed proteins (Supp Figure 4 heat map (B) values >0.2) of low response lines following simvastatin treatment.

Supplemental Table 9: List of proteins used for Heat map analysis of proteins showing differential expression ($|\log_2 FC| > 0.263$) between high and low response hiPSC lines following atorvastatin treatment (Figure 4B) and simvastatin treatment (Supp Figure 4B).

1. Y. E. Pearson *et al.*, A statistical framework for high-content phenotypic profiling using cellular feature distributions. *Commun Biol* **5**, 1409 (2022).

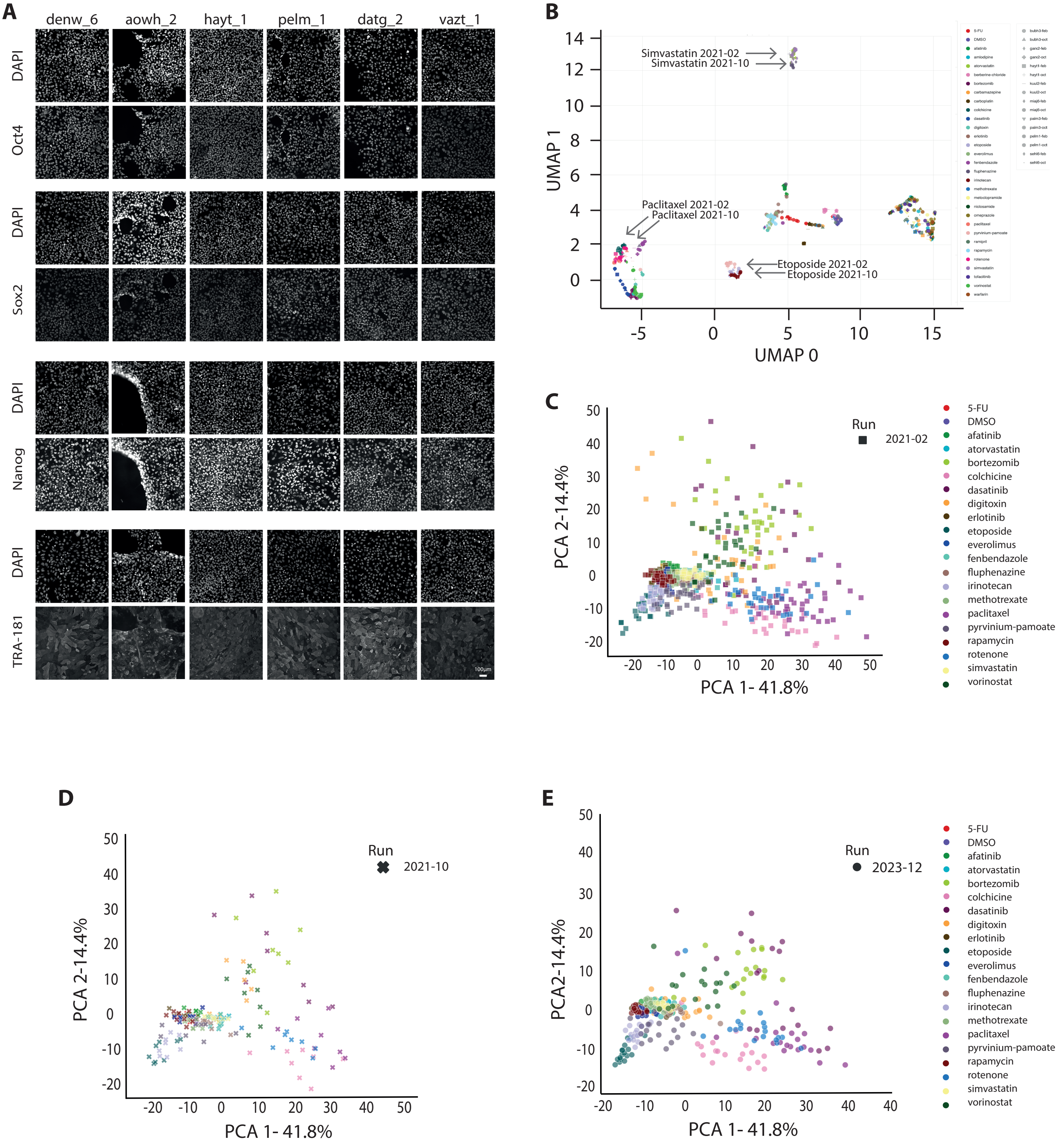


**Supp Figure 1**

A

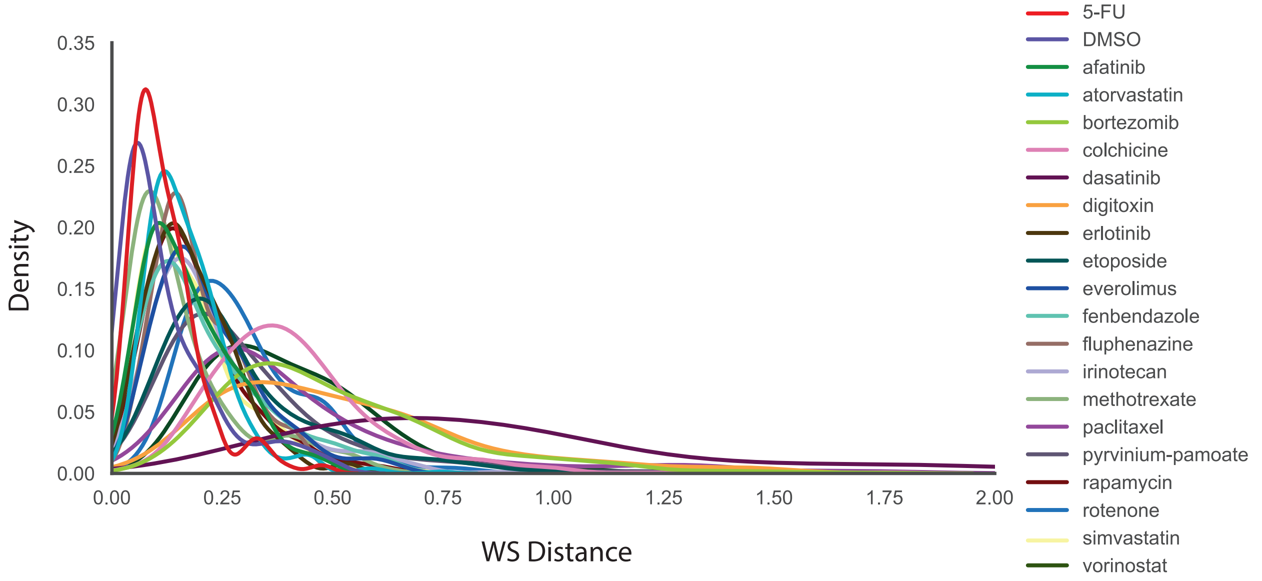


B

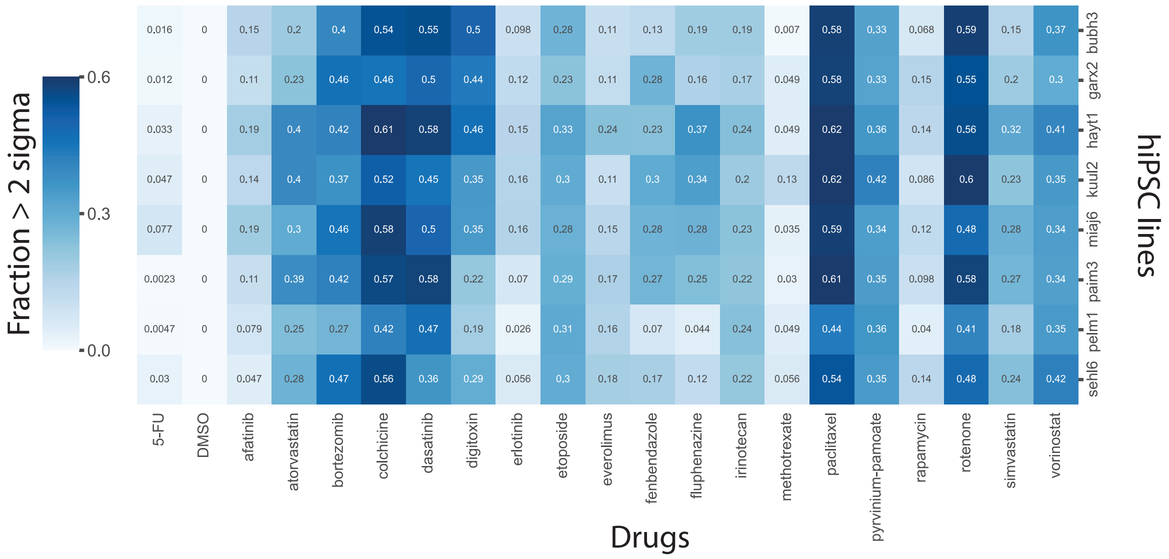


C

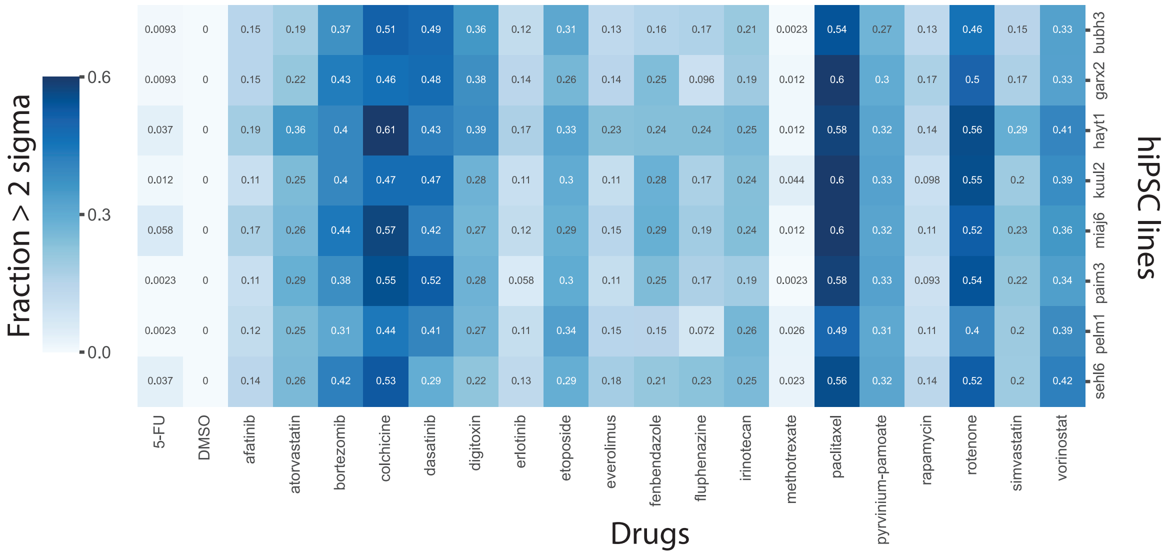


**Supp Figure 2**

A

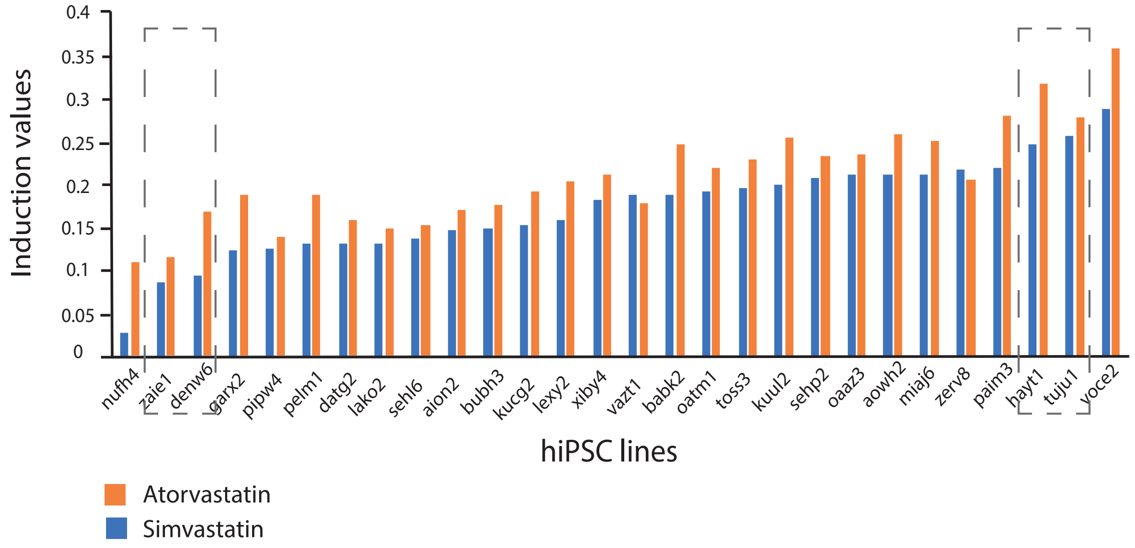


B

Drugs

| hiPSC lines | DMSO | 5-FU | methotrexate | afatinib | erlotinib | fenbendazole | everolimus | simvastatin | rapamycin | fluphenazine | atorvastatin | irinotecan | pyrvinium-pamoate | etoposide | digitoxin | colchicine | rotenone | bortezomib | dasatinib | paclitaxel |
|---|---|---|---|---|---|---|---|---|---|---|---|---|---|---|---|---|---|---|---|---|
| aion2 | 0.00 | 4.83 | 5.46 | 8.87 | 9.20 | 9.95 | 10.04 | 10.52 | 10.79 | 11.62 | 11.97 | 14.48 | 15.15 | 20.27 | 27.65 | 34.65 | 36.77 | 38.69 | 49.15 | 49.23 |
| aowh2 | 0.00 | 8.60 | 7.76 | 16.14 | 24.87 | 12.60 | 19.15 | 18.69 | 18.91 | 17.30 | 22.22 | 28.75 | 38.39 | 53.37 | 62.77 | 61.10 | 41.59 | 53.77 | 52.64 | 63.82 |
| babk2 | 0.00 | 9.80 | 7.40 | 10.25 | 16.56 | 24.51 | 23.32 | 14.84 | 20.95 | 9.92 | 19.89 | 27.54 | 31.67 | 38.66 | 62.68 | 72.68 | 77.59 | 72.19 | 89.28 | 76.14 |
| bubh3 | 0.00 | 4.39 | 6.08 | 10.20 | 8.41 | 9.10 | 10.20 | 9.94 | 9.17 | 9.75 | 13.78 | 17.68 | 19.35 | 25.59 | 51.90 | 50.18 | 40.01 | 48.45 | 48.05 | 54.21 |
| datg2 | 0.00 | 6.18 | 8.10 | 9.45 | 10.96 | 10.51 | 12.31 | 9.95 | 12.01 | 7.85 | 11.44 | 17.44 | 21.76 | 30.05 | 37.87 | 51.16 | 53.36 | 39.60 | 56.47 | 57.97 |
| denw6 | 0.00 | 4.75 | 5.70 | 8.19 | 10.88 | 10.52 | 9.97 | 7.73 | 10.83 | 6.09 | 10.26 | 16.23 | 21.81 | 23.88 | 37.46 | 46.48 | 36.56 | 40.61 | 35.24 | 49.69 |
| garx2 | 0.00 | 3.99 | 7.47 | 8.35 | 11.79 | 13.02 | 11.20 | 9.53 | 10.56 | 7.34 | 10.41 | 16.37 | 17.63 | 22.18 | 31.71 | 36.96 | 36.83 | 50.37 | 45.79 | 47.99 |
| hayt1 | 0.00 | 8.28 | 5.93 | 13.94 | 16.98 | 13.99 | 19.75 | 22.11 | 19.94 | 18.04 | 27.73 | 29.64 | 35.55 | 43.54 | 57.77 | 84.55 | 49.74 | 70.51 | 63.77 | 83.03 |
| kucg2 | 0.00 | 5.62 | 8.87 | 9.68 | 17.11 | 9.12 | 13.81 | 11.39 | 12.04 | 7.40 | 15.59 | 15.48 | 34.61 | 29.62 | 72.83 | 31.49 | 45.27 | 55.08 | 24.59 | 57.22 |
| kuul2 | 0.00 | 7.39 | 6.15 | 9.77 | 15.22 | 17.07 | 14.04 | 15.51 | 14.76 | 16.64 | 20.32 | 25.59 | 21.86 | 31.80 | 47.05 | 66.82 | 64.25 | 58.10 | 73.73 | 76.51 |
| lako2 | 0.00 | 5.60 | 4.15 | 11.11 | 16.30 | 14.53 | 15.40 | 10.26 | 16.16 | 5.77 | 13.65 | 20.77 | 25.50 | 34.15 | 39.40 | 69.31 | 64.79 | 53.65 | 55.08 | 71.88 |
| lexy2 | 0.00 | 5.45 | 5.28 | 6.28 | 9.98 | 13.82 | 9.65 | 9.87 | 9.75 | 8.81 | 13.83 | 15.39 | 28.24 | 20.13 | 45.70 | 45.68 | 53.69 | 44.62 | 52.22 | 67.54 |
| miaj6 | 0.00 | 10.42 | 8.98 | 13.38 | 16.48 | 17.75 | 17.71 | 19.50 | 19.81 | 14.61 | 22.43 | 29.17 | 31.10 | 41.80 | 31.93 | 75.06 | 46.15 | 68.70 | 62.24 | 58.55 |
| nufh4 | 0.00 | 5.93 | 8.55 | 7.35 | 7.56 | 9.61 | 8.86 | 6.93 | 9.74 | 7.58 | 8.45 | 11.10 | 14.41 | 19.16 | 68.08 | 28.14 | 30.83 | 40.47 | 33.36 | 41.84 |
| oaaz3 | 0.00 | 5.46 | 10.52 | 7.54 | 11.36 | 13.27 | 12.75 | 16.39 | 14.14 | 12.64 | 18.71 | 22.76 | 27.66 | 33.61 | 33.61 | 57.09 | 35.58 | 51.29 | 34.11 | 51.46 |
| oatm1 | 0.00 | 7.42 | 5.84 | 12.00 | 14.58 | 12.82 | 14.11 | 14.91 | 13.24 | 10.23 | 17.49 | 26.72 | 23.91 | 32.81 | 45.87 | 67.55 | 46.54 | 49.38 | 59.05 | 66.11 |
| paim3 | 0.00 | 4.45 | 5.99 | 9.01 | 7.17 | 14.68 | 13.66 | 14.71 | 12.62 | 11.37 | 19.73 | 23.14 | 25.60 | 28.73 | 17.03 | 55.78 | 48.91 | 57.82 | 54.43 | 60.19 |
| pelm1 | 0.00 | 4.90 | 5.56 | 8.55 | 5.34 | 7.17 | 11.61 | 11.54 | 9.14 | 6.07 | 14.36 | 18.09 | 29.63 | 25.75 | 21.20 | 30.39 | 27.55 | 39.08 | 27.61 | 30.87 |
| pipw4 | 0.00 | 4.48 | 3.69 | 7.51 | 8.49 | 10.85 | 10.94 | 9.29 | 9.84 | 5.94 | 10.82 | 14.47 | 17.61 | 21.73 | 34.22 | 36.53 | 36.03 | 42.25 | 32.40 | 44.36 |
| sehl6 | 0.00 | 4.48 | 6.42 | 8.45 | 8.49 | 12.44 | 12.48 | 14.22 | 12.41 | 8.89 | 16.47 | 22.09 | 23.40 | 33.59 | 22.69 | 55.08 | 42.05 | 34.91 | 56.77 | 55.53 |
| sehp2 | 0.00 | 9.23 | 7.40 | 11.03 | 12.32 | 15.35 | 19.45 | 18.83 | 18.27 | 11.99 | 22.55 | 30.65 | 29.19 | 49.23 | 42.07 | 74.62 | 51.73 | 65.80 | 65.46 | 84.09 |
| toss3 | 0.00 | 6.92 | 6.94 | 13.29 | 19.63 | 15.46 | 18.40 | 17.07 | 19.45 | 7.92 | 21.72 | 31.62 | 34.47 | 48.19 | 36.03 | 72.35 | 51.55 | 68.82 | 48.68 | 70.28 |
| tuju1 | 0.00 | 6.83 | 10.38 | 12.90 | 18.95 | 14.33 | 13.76 | 22.23 | 16.43 | 20.33 | 24.86 | 32.37 | 38.80 | 54.31 | 29.43 | 69.67 | 48.73 | 59.67 | 39.06 | 64.29 |
| vazt1 | 0.00 | 5.29 | 5.43 | 9.57 | 13.76 | 13.46 | 11.27 | 14.52 | 11.98 | 13.45 | 14.75 | 22.04 | 22.06 | 37.38 | 27.96 | 56.02 | 44.11 | 45.16 | 34.03 | 57.37 |
| voce2 | 0.00 | 6.98 | 6.23 | 15.60 | 14.75 | 20.33 | 13.21 | 22.71 | 13.24 | 25.55 | 31.48 | 25.33 | 28.45 | 41.22 | 54.92 | 63.16 | 66.95 | 72.71 | 73.52 | 67.32 |
| xiby4 | 0.00 | 6.91 | 7.22 | 10.43 | 21.41 | 10.19 | 15.19 | 11.94 | 17.80 | 5.89 | 17.44 | 27.60 | 31.68 | 39.34 | 47.42 | 71.04 | 45.36 | 58.93 | 41.55 | 69.43 |
| zaie1 | 0.00 | 3.72 | 5.98 | 8.97 | 5.28 | 8.84 | 8.34 | 8.90 | 9.51 | 7.85 | 12.05 | 15.19 | 17.83 | 23.92 | 21.02 | 38.30 | 29.48 | 33.02 | 39.07 | 40.13 |
| zerv8 | 0.00 | 4.98 | 5.97 | 12.08 | 12.07 | 12.64 | 10.58 | 16.97 | 11.06 | 15.63 | 18.86 | 16.79 | 21.66 | 29.50 | 35.28 | 50.40 | 45.36 | 44.58 | 33.44 | 53.52 |
| | Sigma | 1.81 | 1.65 | 2.51 | 4.91 | 3.72 | 3.76 | 4.57 | 3.81 | 4.99 | 5.63 | 6.25 | 6.90 | 10.18 | 14.84 | 15.88 | 11.73 | 11.88 | 15.54 | 12.95 |
| | Mean | 6.19 | 6.77 | 10.35 | 13.07 | 13.14 | 13.61 | 13.96 | 13.73 | 11.16 | 17.26 | 21.95 | 26.04 | 33.34 | 40.84 | 55.44 | 46.33 | 52.08 | 49.31 | 59.66 |
| | CoV | 0.29 | 0.24 | 0.24 | 0.38 | 0.28 | 0.28 | 0.33 | 0.28 | 0.45 | 0.33 | 0.28 | 0.26 | 0.31 | 0.36 | 0.29 | 0.25 | 0.23 | 0.32 | 0.22 |

**Supp Figure 3**

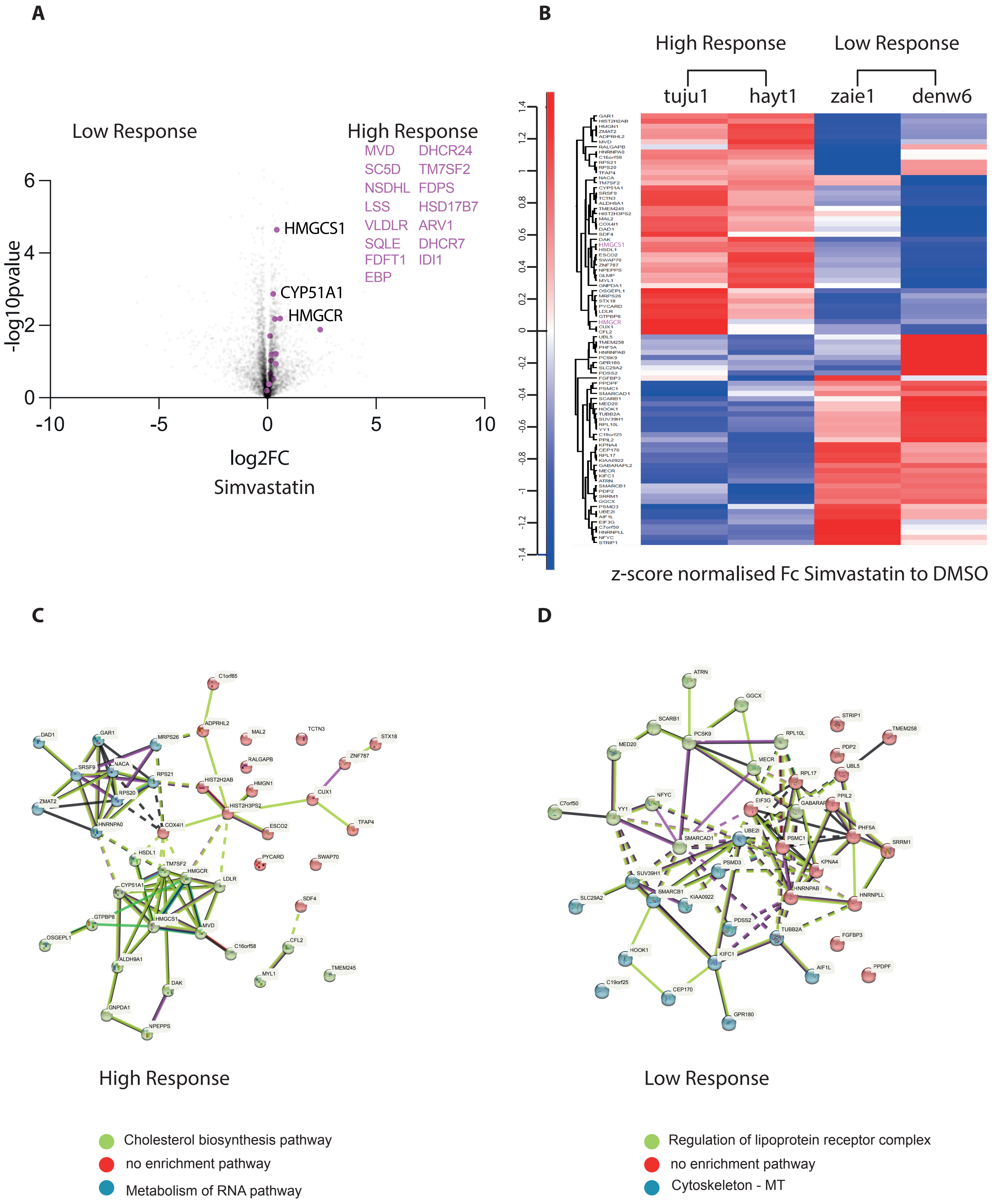


**Supp Figure 4**

**A**

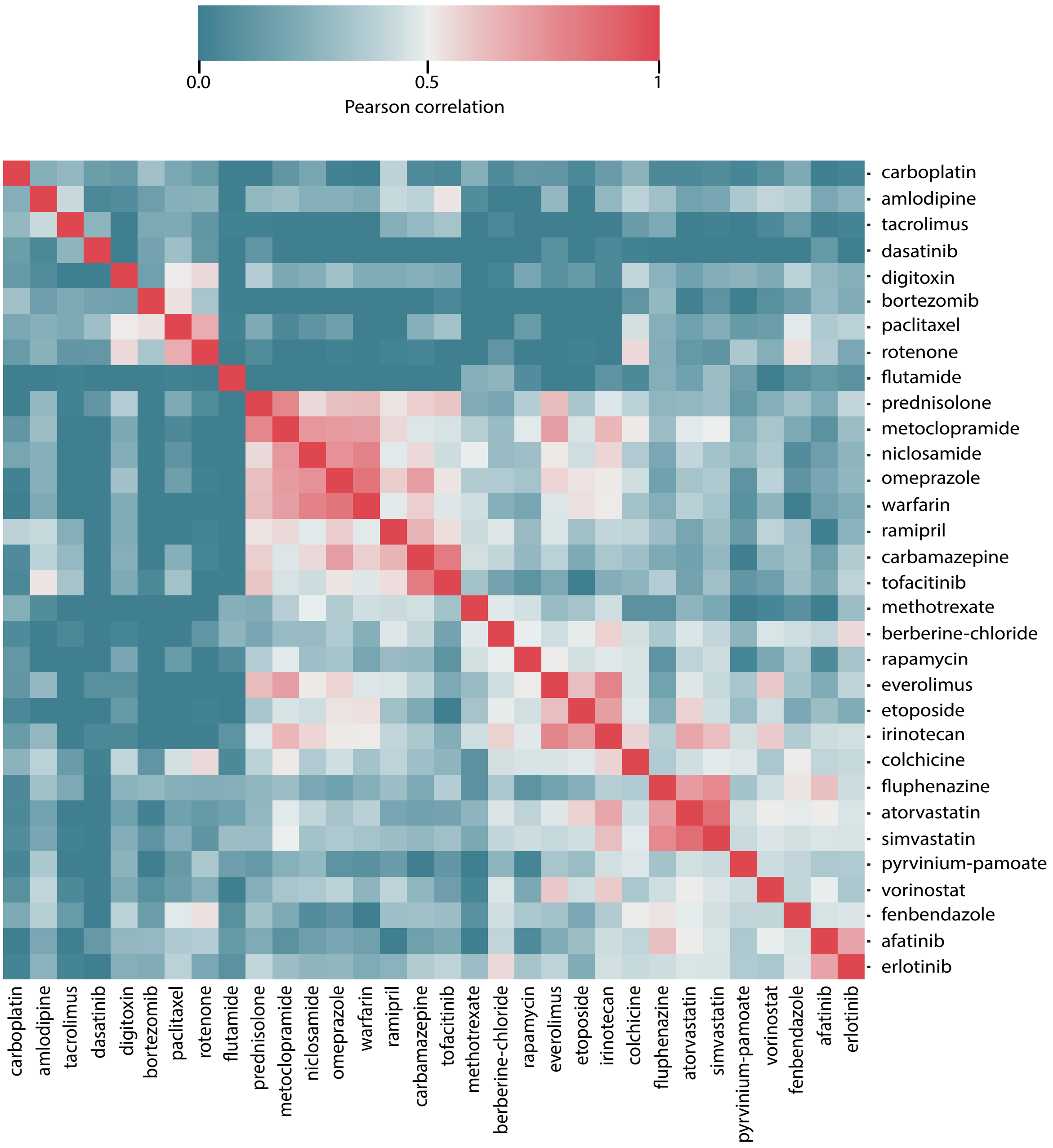


**Supp Figure 5**

**Supplemental Table 1: List of hiPSC lines used in this study**

| Cell ID | Donor Description | Run #2021-02 | Run #2021-10 | Run #2023-12 |
|---|---|---|---|---|
| aion2 | White British, Male | ✓ | | |
| aowh2 | White British, Female | ✓ | | |
| babk2 | White British, Female | ✓ | | |
| bubh3 | White British, Female | ✓ | | |
| burb1 | White British, Male | | ✓ | ✓ |
| datg2 | White British, Female | ✓ | | |
| denw6 | White British, Male | ✓ | | ✓ |
| deyz2 | White British, Female | | | ✓ |
| fiaj1 | White British, Male | | | ✓ |
| garx2 | White British, Female | ✓ | ✓ | |
| hayt1 | White Other, Male | ✓ | ✓ | ✓ |
| hiaf1 | White British, Male | | | ✓ |
| kucg2 | White British, Male | ✓ | | |
| kuul2 | White British, Male | ✓ | ✓ | |
| lako2 | White British, Female | ✓ | | |
| lexy2 | White British, Female | ✓ | | |
| melw1 | White British, Male | | | ✓ |
| miaj6 | White British, Male | ✓ | ✓ | |
| nufh4 | White British, Female | ✓ | | |
| oaaz3 | White British, Male | ✓ | | |
| oatm1 | White British, Male | ✓ | | |
| paim3 | White British, Male | ✓ | ✓ | |
| pelm1 | White British, Female | ✓ | ✓ | |
| pipw4 | White British, Male | ✓ | | |
| podx2 | White British, Female | | | ✓ |
| puhk2 | White British, Female | | | ✓ |
| romx1 | White British, Male | | | ✓ |
| sebn3 | White British, Female | | | ✓ |
| sehl6 | White British, Female | ✓ | ✓ | |
| sehp2 | White Other, Female | ✓ | | |
| sohd3 | White British, Female | | | ✓ |
| tert1 | White British, Male | | | ✓ |
| toss3 | White British, Male | ✓ | | |
| tuju1 | White British, Female | ✓ | | ✓ |
| vaka5 | White British, Female | | | ✓ |
| vazt1 | White British, Male | ✓ | | |
| voce2 | White British, Male | ✓ | | |
| wibj2 | White British, Female | | | ✓ |
| xiby4 | White British, Female | ✓ | | |
| zaie1 | White British, Female | ✓ | | |
| zapk3 | White British, Male | | | ✓ |
| zerv8 | White British, Female | ✓ | | |
| zoxy3 | White British, Female | | | ✓ |

**Supplemental Table 2: List of drugs and concentrations used in this study**

| Drug | Disease target | Concentration |
|---|---|---|
| **DMSO (control)** | *Control* | 5 µM |
| **5-FU** | Cancer drug | 5 µM |
| **Abacavir** | HIV reverse transcriptase inhibitor | 5 µM |
| **Afatinib** | Cancer drug (breast) | 5 µM |
| **Amlodipine** | Heart disease | 5 µM |
| **Atorvastatin** | Heart disease (coronary) | 5 µM |
| **Azathioprine** | Immunosuppressant | 5 µM |
| **Berberine Chloride** | Transcription/mitochondrial function | 2.68 µM |
| **Bortezomib** | Cancer drug | 5 µM |
| **Bosentan** | Heart disease (Pulmonary hypertension) | 5 µM |
| **Carbamazepine** | Epilepsy | 5 µM |
| **Carboplatin** | Cancer drug (neoplasm/carcinoma) | 5 µM |
| **Cyclophosphamide Monohydrate** | Cancer drug (breast neoplasm) | 5 µM |
| **Dasatinib** | Cancer drug | 5 µM |
| **Dexamethasone** | Lung condition | 5 µM |
| **Digitoxin** | Cancer drug | 5 µM |
| **Erlotinib** | Cancer drug (breast) | 5 µM |
| **Ethacrynic Acid** | Cancer drug, ALS, Parkinson’s | 5 µM |
| **Etoposide** | Cancer drug (topoisomerase inhibitor) | 1.7 µM |
| **Everolimus** | Cancer drug (breast) | 5 µM |
| **Fenbendazole** | Parasitic infections | 3.34 µM |
| **Fluphenazine** | Antipsychotic | 5 µM |
| **Flutamide** | Cancer drug (prostate) | 5 µM |
| **Gefitinib** | Cancer drug (breast) | 5 µM |
| **Hydrocortisone** | Steroid | 5 µM |
| **Imatinib** | Cancer drug | 5 µM |
| **Irinotecan** | Cancer drug (topoisomerase inhibitor) | 5 µM |
| **Lansoprazole** | Stomach ulcer | 5 µM |
| **Metformin** | Diabetes | 5 µM |
| **Methotrexate** | Cancer drug (lymphoma) | 5 µM |
| **Metoclopramide** | Nausea/vomiting | 5 µM |
| **Milrinone** | Heart disease (heart failure) | 5 µM |
| **Niclosamide** | Cancer/bacterial/viral infections | 5 µM |
| **Omeprazole** | Stomach ulcer | 5 µM |
| **Paclitaxel** | Cancer drug (breast) | 5 µM |
| **Paroxetine** | Depressive disorder | 5 µM |
| **Prednisolone** | Carotid stenosis | 5 µM |
| **Procaine** | Anaesthesia | 5 µM |
| **Pyrvinium Pamoate** | Antihelminthic drug - anticancer | 5 µM |

| | | |
|---|---|---|
| **Ramipril** | Heart disease | 5 μM |
| **Rapamycin** | mTOR inhibitor | 5 μM |
| **Rosiglitazone** | Diabetes | 5 μM |
| **Rotenone** | Pesticide | 5 μM |
| **Salbutamol** | Bronchodilator | 5 μM |
| **Simvastatin** | Coronary disease | 5 μM |
| **Sorafenib** | Cancer drug (breast) | 5 μM |
| **Tacrolimus** | Immunosuppressant | 5 μM |
| **Tofacitinib** | Arthritis | 5 μM |
| **Voriconazole** | Anti-fungal | 5 μM |
| **Vorinostat** | HDAC inhibitor | 5 μM |
| **Warfarin** | Heart disease | 5 μM |

**Supplemental Table 3: GO enrichment analysis of top 5% proteins showing increased expression in all four hiPSC lines used for proteomic analysis following atorvastatin treatment.**

| GO term | Description | p-value | FDR q-value | Enrichment |
|---|---|---|---|---|
| GO:0016126 | sterol biosynthetic process | 1.71E-20 | 1.37E-16 | 16.74 |
| GO:0008610 | lipid biosynthetic process | 5.99E-20 | 2.39E-16 | 8.06 |
| GO:0006695 | cholesterol biosynthetic process | 3.56E-19 | 9.47E-16 | 16.59 |
| GO:1902653 | secondary alcohol biosynthetic process | 3.56E-19 | 7.11E-16 | 16.59 |
| GO:0016125 | sterol metabolic process | 9.72E-19 | 1.55E-15 | 12.47 |
| GO:0006694 | steroid biosynthetic process | 6.08E-18 | 8.10E-15 | 12.66 |
| GO:0008202 | steroid metabolic process | 1.00E-17 | 1.14E-14 | 9.85 |
| GO1902652 | secondary alcohol metabolic process | 1.54E-17 | 1.54E-14 | 12.23 |
| GO:000820 | cholesterol metabolic process | 1.54E-17 | 1.37E-14 | 12.23 |
| GO:0006629 | lipid metabolic process | 1.94E-16 | 1.55E-13 | 4.96 |
| Go:0046165 | alcohol biosynthetic process | 7.83E-15 | 5.69E-12 | 11.25 |
| GO:1901617 | organic hydroxy compound biosynthetic process | 2.03E-14 | 1.35E-11 | 9.85 |
| GO:0019216 | regulation of lipid metabolic process | 2.55E-14 | 1.57E-11 | 7.72 |
| GO:0046890 | regulation of lipid biosynthetic process | 6.87E-14 | 3.92E-11 | 10.17 |
| GO:0050810 | regulation of steroid biosynthetic process | 8.11E-13 | 4.32E-10 | 12.19 |
| GO:0006066 | alcohol metabolic process | 1.02E-12 | 5.07E-10 | 7.54 |
| GO:1901615 | organic hydroxy compound metabolic process | 1.40E-12 | 6.59E-10 | 6.46 |
| GO:1902930 | regulation of alcohol biosynthetic process | 1.85E-12 | 8.19E-10 | 13.13 |
| GO:0045540 | regulation of cholesterol biosynthetic process | 1.85E-12 | 7.76E-10 | 13.13 |
| GO:0090181 | regulation of cholesterol metabolic process | 1.85E-12 | 7.37E-10 | 13.13 |
| GO:0106118 | regulation of sterol biosynthetic process | 1.85E-12 | 7.02E-10 | 13.13 |

**Supplemental Table 4: GO enrichment analysis of top 5% proteins showing increased expression in all four hiPSC lines used for proteomic analysis following simvastatin treatment.**

| GO Term | Description | P- value | FDR q-value | Enrichment |
|---|---|---|---|---|
| GO:0016125 | sterol metabolic process | 6.66E-16 | 7.50E-13 | 10.483 |
| GO:1902652 | secondary alcohol metabolic process | 6.66E-16 | 7.50E-13 | 10.483 |
| GO:0008203 | cholesterol metabolic process | 1.02E-14 | 5.67E-12 | 10.207 |
| GO:0008202 | steroid metabolic process | 1.49E-14 | 5.67E-12 | 8.325 |
| GO:0016126 | sterol biosynthetic process | 1.51E-14 | 5.67E-12 | 11.1 |
| GO:1902653 | secondary alcohol biosynthetic process | 1.51E-14 | 5.67E-12 | 11.1 |
| GO:0090181 | regulation of cholesterol metabolic process | 3.34E-14 | 1.08E-11 | 13.361 |
| GO:0019218 | regulation of steroid metabolic process | 1.06E-13 | 2.98E-11 | 11.261 |
| GO:0008610 | Lipid biosynthetic process | 1.30E-13 | 3.26E-11 | 5.9769 |
| GO:0006694 | steroid biosynthetic process | 1.54E-13 | 3.46E-11 | 8.9697 |
| GO:0006695 | cholesterol biosynthetic process | 2.45E-13 | 5.02E-11 | 10.792 |
| GO:0006066 | alcohol metabolic process | 3.35E-13 | 6.29E-11 | 7.8625 |
| GO:1901615 | organic hydroxy compound metabolic process | 4.83E-13 | 8.37E-11 | 7.0851 |
| GO:0045540 | regulation of cholesterol biosynthetic process | 6.13E-13 | 8.63E-11 | 13.059 |
| GO:0106118 | regulation of sterol biosynthetic process | 6.13E-13 | 8.63E-11 | 13.059 |
| GO:1902930 | regulation of alcohol biosynthetic process | 6.13E-13 | 8.63E-11 | 13.059 |
| GO:0050810 | regulation of steroid biosynthetic process | 7.13E-13 | 9.45E-11 | 11.452 |
| GO:1901617 | organic hydroxy compound biosynthetic process | 8.70E-13 | 1.09E-10 | 8.2222 |
| GO:0046165 | alcohol biosynthetic process | 1.13E-12 | 1.34E-10 | 8.9516 |
| GO:0046890 | regulation of lipid biosynthetic process | 8.12E-12 | 9.15E-10 | 8.931 |
| GO:0044283 | small molecule biosynthetic process | 3.45E-11 | 3.70E-09 | 5.3258 |
| GO:0006629 | lipid metabolic process | 4.54E-11 | 4.65E-09 | 4.0913 |
| GO:0062012 | regulation of small molecule metabolic process | 4.81E-10 | 4.71E-08 | 7 |
| GO:0019216 | regulation of lipid metabolic process | 1.64E-09 | 1.54E-07 | 6.475 |

**Supplemental Table 5: GO Enrichment Analysis of gene sets/pathways and their associated p-values for highly expressed proteins (Fig 4 heat map (B) values >0.2) of High response lines following atorvastatin treatment.**

| Gene set | Description | Ratio | p-value | FDR |
|---|---|---|---|---|
| GO:0090181 | Regulation of cholesterol metabolic process | 19.427 | 4.6062e-7 | 0.0018237 |
| GO:0008203 | cholesterol metabolic process | 12.592 | 0.0000010172 | 0.0018237 |
| GO:0046890 | Regulation of lipid biosynthetic process | 12.395 | 0.0000011345 | 0.0018237 |
| GO:1902652 | Secondary alcohol metabolic process | 12.204 | 0.0000012630 | 0.0018237 |
| GO:0006695 | cholesterol biosynthetic process | 15.454 | 0.0000018865 | 0.0018237 |
| GO:0016125 | Sterol metabolic process | 11.497 | 0.0000019057 | 0.0018237 |
| GO:1902653 | Secondary alcohol biosynthetic process | 15.110 | 0.0000021623 | 0.0018237 |
| GO:0016126 | Sterol biosynthetic process | 14.467 | 0.0000028127 | 0.0018452 |
| GO:0019218 | Regulation of steroid metabolic process | 14.467 | 0.0000028127 | 0.0018452 |
| GO:0045540 | Regulation of cholesterol biosynthetic process | 18.278 | 0.0000062092 | 0.0030549 |

**Supplemental Table 6: GO Enrichment Analysis of gene sets/pathways and their associated p-values for highly expressed proteins (Fig 4 heat map (B) values >0.2) of low response lines following atorvastatin treatment.**

| Gene set | Description | Ratio | p-value | FDR |
|---|---|---|---|---|
| GO:0043687 | Post-translational protein modification | 6.8556 | 3.6169e-7 | 0.0021354 |
| GO:0006814 | Sodium ion transport | 9.7013 | 0.00014502 | 0.39546 |
| GO:0043062 | Extracellular structure organization | 5.6680 | 0.00020094 | 0.39546 |
| GO:0035752 | Sodium ion transmembrane trasnport | 11.752 | 0.00033453 | 0.49377 |
| GO:0098869 | Cellular oxidant detoxification | 9.3485 | 0.00081295 | 0.51703 |
| GO:0034368 | Protein-lipid complex remodeling | 41.133 | 0.00091104 | 0.51703 |
| GO:0034369 | Plasma lipoprotein particle remodeling | 41.133 | 0.00091104 | 0.51703 |
| GO:0034375 | High-density lipoprotein particle remodeling | 41.133 | 0.00091104 | 0.51703 |
| GO:2000644 | Regulation of receptor catabolic process | 41.133 | 0.00091104 | 0.51703 |
| GO:1905906 | Regulation of amyloid fibril formation | 41.133 | 0.00091104 | 0.51703 |

**Supplemental Table 7: GO Enrichment Analysis of gene sets/pathways and their associated p-values for highly expressed proteins (Supp Fig 2 heat map (B) values >0.2) of High response lines following simvastatin treatment.**

| Gene set | Description | Ratio | p-value | FDR |
|---|---|---|---|---|
| GO:0090181 | Regulation of cholesterol metabolic process | 25.728 | 8.0828e-8 | 0.00047721 |
| GO:0019218 | Regulation of steroid metabolic process | 19.159 | 5.0470e-7 | 0.0014899 |
| GO:0045540 | Regulation of cholesterol biosynthetic process | 24.207 | 0.0000014877 | 0.0017567 |
| GO:0106118 | Regulation of sterol biosynthetic process | 24.207 | 0.0000014877 | 0.0017567 |
| GO:1902930 | Regulation of alcohol biosynthetic process | 24.207 | 0.0000014877 | 0.0017567 |
| GO:0008203 | Cholesterol metabolic process | 14.293 | 0.0000029569 | 0.0026274 |
| GO:0046890 | Regulation of lipid biosynthetic process | 14.070 | 0.0000032471 | 0.0026274 |
| GO:1902652 | Secondary alcohol metabolic process | 13.854 | 0.0000035601 | 0.0026274 |
| GO:0050810 | Regulation of steroid biosynthetic process | 19.241 | 0.0000048505 | 0.0029923 |
| GO:0016125 | Sterol metabolic process | 13.051 | 0.0000050683 | 0.0029923 |

**Supplemental Table 8: GO Enrichment Analysis of gene sets/pathways and their associated p-values for highly expressed proteins (Supp Fig 2 heat map (B) values >0.2) of low response lines following simvastatin treatment.**

| Gene set | Description | Ratio | p-value | FDR |
|---|---|---|---|---|
| GO:0036123 | Histone H3-K9 dimethylation | 58.453 | 0.00045016 | 1 |
| GO:0018027 | Peptidyl-lysine dimethylation | 29.226 | 0.0019824 | 1 |
| GO:0036124 | Histone H3-K9 tri methylation | 26.569 | 0.0024125 | 1 |
| GO:0051567 | Histone H3-K9 methylation | 15.382 | 0.0072464 | 1 |
| GO:0042632 | Cholesterol homeostasis | 13.917 | 0.0088228 | 1 |
| GO:0008203 | Cholesterol metabolic process | 6.9586 | 0.0088508 | 1 |
| GO:1902652 | Secondary alcohol metabolic process | 6.7445 | 0.0096445 | 1 |
| GO:0055092 | Sterol homeostasis | 13.285 | 0.0096635 | 1 |
| GO:0034383 | Low-density lipoprotein particle clearance | 12.707 | 0.010538 | 1 |
| GO:0016125 | Sterol metabolic process | 6.3535 | 0.0011354 | 1 |
